# Linking Anthocyanin Diversity, Hue, And Genetics In Purple Corn

**DOI:** 10.1101/2020.05.15.098756

**Authors:** Laura A. Chatham, John A. Juvik

## Abstract

While maize with anthocyanin-rich pericarp (purple corn) is rising in popularity as a source of natural colorant for foods and beverages, information on color range and stability—factors associated with anthocyanin decorations and compositional profiles—are currently limited. Using the natural anthocyanin diversity present in a purple corn landrace, Apache Red, we generated a population with variable flavonoid profiles—flavonol-anthocyanin condensed forms (0-83%), acylated anthocyanins (2-72%), pelargonidin-derived anthocyanins (5-99%), and C-glycosyl flavone co-pigments up to 1904 µg/g—all of which contributed in part to the absorbance profile, used here as a proxy for hue. This variability offers targets of selection for breeders looking to expand both stability and the available range of colors that can be sourced from purple corn. With genotyping-by-sequencing of this population we mapped these anthocyanin profile traits. Major QTL for anthocyanin type were found near loci previously identified only in aleurone-pigmented maize varieties [*Purple aleurone1* (*Pr1*) and *Anthocyanin acyltransferase1* (*Aat1*)]. A QTL near *P1* (*Pericarp color1*) was found for both flavone content and flavanol-anthocyanin condensed forms. A significant QTL associated with peonidin-derived anthocyanins near a candidate S-adenosylmethionine-dependent methyltransferase was also identified, warranting further investigation. This population represents the most anthocyanin diverse pericarp-pigmented maize variety characterized to date. Moreover, the candidates identified here will serve as branching points for future research studying the genetic and molecular processes determining anthocyanin profile in pericarp.

## Introduction

Purple corn could provide an economical and natural source of colorants, but as demand rises and the desired application ranges broaden, new challenges are likely to emerge. Matching the range of possible colors, stability, and price that had been attained previously with artificial colorants will likely require additional development through breeding. A major breeding objective for purple corn is to expand its food and beverage applicability by increasing the range of available colors and shelf life. However, these goals must be accomplished while maintaining grain yield and other agronomic traits.

Understanding the genetics underlying anthocyanin biosynthesis in purple corn could expedite the breeding of new cultivars. Kernel anthocyanin content has long been studied in maize due to its easily observable phenotype, but most research to date has focused on kernels with pigmented aleurone, the outermost, single cell layer of the triploid endosperm. However, purple corn contains anthocyanin in the pericarp, the maternal, outermost layer of the kernel. Compared to aleurone, pericarp pigmented lines comprise a relatively small portion of the available germplasm and thus have limited variability. Despite this disadvantage, lines with anthocyanin-pigmented pericarp often have concentrations an order of magnitude greater than aleurone lines^1^. Anthocyanin extraction may also be more efficient from pericarp lines^2^. Moreover, established milling processes separating pericarp could provide valuable pigment-rich pericarp fractions and non-pigmented grain and germ fractions to be used for the production of food, feed, or fuel, creating a value-added product^3^.

Anthocyanins in purple corn consist primarily of the glucosides, and malonyl and dimalonyl glucosides of the anthocyanidins cyanidin, pelargonidin, and peonidin (Supplementary Figure 1). Color is largely influenced by anthocyanidin type, shifting from orange/red to pink/red as the number of hydroxyl and methoxy groups on the B ring increases^4^. However, anthocyanidin type alone cannot account for the diversity in colors observed. Color stability and extract shelf life have been associated with the proportion of acylated anthocyanins^5^. Another category of anthocyanin types is the flavonol-anthocyanin condensed forms, heterodimers consisting of a flavan-3-ol covalently linked to an anthocyanin, but these have only been found in significant quantities in pericarp pigmented varieties^1^. Condensed forms have also been associated with both altered hues and stability^6,7^.

The anthocyanin biosynthesis pathway in maize is well studied and most structural genes involved, with the exception of those controlling the formation of peonidin-based anthocyanins and flavonol-anthocyanin condensed forms, have been characterized^8,9^. However, these structural genes were primarily identified in aleurone pigmented varieties. While the pathway is well-conserved among plants and most structural genes are assumed to operate similarly in aleurone and pericarp, empirical evidence of this is lacking. The genes and mechanisms controlling anthocyanidin type (e.g. pelargonidin, cyanidin, peonidin) and anthocyanin decorations such as acylation and condensation with flavanols are of interest in purple corn because of their potential to influence the color and stability of extracts. *Pr1* (*Red aleurone1*) is known to control the ratio of cyanidin and pelargonidin in aleurone^9^, and more recently, an anthocyanin acyltransferase (*Aat1*) was identified in an aleurone pigmented population with reduced proportions of acylated anthocyanins in some individuals^10^. This work seeks to confirm whether these same genes function in pericarp and to identify loci associated with the biosynthesis of peonidin and condensed forms.

Another factor controlling purple corn extract color and stability is copigmentation. The addition of maize C-glycosyl flavone-rich extract to maize anthocyanin-rich extract can significantly alter hue and stability^11^. Flavone and anthocyanin co-occurrence in the same lines and its effect on extract hue has not been studied or reported previously. *Pericarp color1* (*P1*) is known to regulate the biosynthesis of flavones^12^ but factors controlling the relative ratios of anthocyanin to flavone within a variety are not known.

Color plays a significant role in a consumer’s perception of food and beverage identity, flavor, and quality^13^, making hue an important consideration in the formulation of a natural colorant. Even subtle differences in product color could have a large impact on consumer acceptance, and thus understanding how various anthocyanin and flavonoid chemistries lead to differences in hue will be important for optimizing color in food and beverage applications. Moreover, linking chemistries and hue to their underlying genetics may aid in breeding purple corn lines for increased color range. Here we seek to explore the most prominent factors contributing to purple corn anthocyanin extract color and to identify the candidate loci responsible.

## Materials and Methods

### Plant materials

Seeds of a purple corn landrace, Apache Red (AR), were purchased from Siskiyou Seeds (Williams, OR)(S_0_ population) and grown out to produce S_1_ seeds. S_1_ seeds were analyzed for anthocyanin content using HPLC and seven lines containing anthocyanins were advanced while non-anthocyanin containing progeny were discarded. Selected anthocyanin-rich S_1_ ears along with a sample of S_0_ seeds from the original AR source were grown and self-pollinated, producing 181 S_2_s and 8 new S_1_s, which were also analyzed using HPLC. The available S_2_ lines were genotyped as described below. A selection (9 lines) of highly pigmented S_2_ seeds were grown in a winter nursery to produce 86 S_3_s. The following season, a selection of anthocyanin containing S_1_, S_2_, and S_3_ lines spanning the available anthocyanin diversity in our AR lines were planted and S_2_, S_3_, and S_4_ seed was recovered. Of these, anthocyanin-rich lines were selected from each S_1_, S_2_, and S_3_ parent for genotyping and phenotyping (1148 lines, 103 S_2_s, 730 S_3_s, and 315 S_4_s). To this, 69 samples containing no or minimal anthocyanin content and representing the original 7 S_1_ families were added and genotyped and phenotyped to supplement the data set. A schematic showing how the population was created can be found in Supplementary Figure 2.

### Extractions

The anthocyanin extraction method was based on previously published protocols and findings regarding the efficiency of anthocyanin extraction protocols^1,14^. Briefly, a subset of kernels from each line was ground to a fine powder using a coffee grinder and 1 g of powder was weighed out for extraction. Samples were extracted in 5 ml of aqueous 2% formic acid for 1 h at 50 °C with constant agitation. Extracts were centrifuged and filtered through a 25 mm, 0.45 µm Millex LCR PTFE syringe filter (Millipore, Billerica, MA) prior to HPLC analysis and optical measurement with UV-visible spectroscopy.

### HPLC analysis

Anthocyanin and flavone content was measured using an Agilent 1100 series high-performance liquid chromatography (HPLC) system and diode array detector (DAD) (Santa Clara, CA) in a similar manner to previous reports^14^. The stationary phase consisted of a Poroshell 120 SB-C18 100 mm x 4.6 mm, 2.7 µm column (Agilent), and the mobile phase was a two-solvent mixture, with 2% formic acid in water as solvent A, and acetonitrile as solvent B. The protocol began at 93% solvent A and 7% solvent B, and solvent B increased linearly to 18% over 30 min. Column temperature was kept at 30 °C, an injection volume of 20 µl was used, and absorbance was measured at 520 for anthocyanin content and 340 nm for flavone content. ChemStation (rev A10.02) software (Agilent) was used for integration and anthocyanins were quantified using standard curves of cyanidin, pelargonidin, and peonidin 3-glucosides, while flavones were quantified using *C-*hexosyl apigenin compounds (vitexin and isovitexin) (Extrasynthese, Genay, France). Identities were confirmed by mass spectrometry as described previously^14^. To ensure consistency between batches of samples and retention times, an AR mixture was created by combining ground powder from multiple AR lines, a sample of which was extracted and run on HPLC with each batch of samples.

### UV-Visible spectroscopy

Absorbance was recorded between 380 and 780nm on a Synergy 2 multi-well plate reader (Biotek, Winooski, VT) using a 96-well plate. Each extract was run in triplicate and the maximum absorbance (Abs_max_) and the wavelength at Abs_max_ (λ_max_) were recorded.

### Statistical analysis

Anthocyanin content, flavone content, and all other factors used in analyzing the relationship between various anthocyanin species and hue were normalized before plotting, and all statistics were calculated using R.

### GBS library construction and sequencing

Fourteen S_2_, S_3_, and S_4_ seeds of each line were sown in 11 x 21 cell propagation trays (Proptek, Watsonville, CA) filled with a soilless seedling mix (Sunshine Redi-Earth Plug & Seedling; SunGro, Agawam, MA) and grown in a greenhouse at the University of Illinois Plant Care Facility. Once seedlings had emerged, tissue of a similar size and proportion was harvested from at least seven seedlings, depending on germination rate. Samples were frozen to -20 °C, lyophilized, and ground in 15ml conical tubes with stainless steel ball bearings using an automated tissue homogenizer (SPEX SamplePrep, Metuchen, NJ). A 96-well optimized CTAB DNA extraction protocol was used for DNA isolation, and concentrations were determined using the Quant-iT PicoGreen dsDNA Assay Kit (Thermo Fisher Scientific, Waltham, MA). GBS libraries were constructed as described previously using a double restriction enzyme digest with *HinP1I* and *PstI*^15,16^. All lines were multiplexed into three lanes and sequenced using the Illumina HiSeq 4000 system with single-end 100 nucleotide reads at the Roy J. Carver Biotechnology Center at the University of Illinois in Urbana, IL.

### SNP discovery and association mapping

SNP discovery was carried out using the TASSEL 5.0 GBS v2 pipeline^17^. Reads were aligned to the *Zea Mays* B73 reference genome version 4^18^ using Bowtie 2^19^, and minor allele frequency was set to 0.05. This generated 149,342 SNP sites with an average depth of 12 and average proportion covered of 0.65. With imputation, performed using Beagle 5.0^20^, and filtering, a total of 41,986 nonredundant SNPs per sample were generated. Association mapping was performed using GAPIT (Genome Association and Prediction Integrated Tool) in R^21^ for several phenotypic traits. Seven principle components were used to adequately account for population structure in addition to the kinship matrix provided by GAPIT.

### Methyltransferase candidates

The amino acid sequence of *VvAOMT1*, an S-adenosyl-L-methionine-dependent methyltransferase, with 3-*O-*methyltransferase activity^22^ was blasted against the maize genome to identify homologous candidates in maize. These candidates along with known caffeoyl-co A *O*-methyltransferases (CCoAOMTs), anthocyanin *O-*methyltransferase (AOMTs) and caffeic acid *O-*methyltransferases (CaOMTs) gathered from NCBI. Candidate sequence selection was based in part on phylograms from Provenzano et al. ^23^ and Du et al.^24^. Sequences were aligned in R using the DECIPHER^25^ package, and a phylogram was created using the phangorn package^26^. Alignment of leucoanthocyanidin reductases and anthocyanidin reductase was performed in a similar manner using the same R packages.

## Results and Discussion

### Anthocyanin diversity

Previously we showed that Apache Red (AR) contains a variety of anthocyanin profiles with some containing mostly pelargonidin-derived anthocyanins, a profile not previously seen in anthocyanin-rich pericarp lines^1,14^. We also identified AR lines with high concentrations of C-glycosyl flavones, which copigment with anthocyanins to alter color intensity, hue, and stability^27^. Given the diversity and unique qualities present in this population, we self-pollinated lines of the original landrace to extract as much phenotypic diversity from the population as possible. The wavelength at maximum absorption (λ_Max_) of anthocyanin lines ranged from 497nm to 521nm. As a proxy for hue, this range suggests good variability in extract colors among the samples. Anthocyanin and C-glycosyl flavones were also highly concentrated compared to previous reports (1598 µg/g whole corn and 1904 µg/g, respectively, Table 1)^1,14^. Many lines were abundant in both, with flavone:anthocyanin ratios as high as 11:1.

**Table 1:** Diversity in anthocyanin content of 1152 Apache Red lines

| Category | Range |
| --- | --- |
| Acylated anthocyanins | 2% - 72% |
| Malonylated | 0% - 43% |
| Dimalonylated | 0% - 57% |
| Pelargonidin-derived anthocyanins | 5% - 99% |
| Cyanidin-derived anthocyanins | 0% - 86% |
| Peonidin-derived anthocyanins | 0% - 50% |
| Flavanol-anthocyanin condensed forms | 0% - 83% |
| Total flavone content | 0-1904 µg/g |
| Total anthocyanin content | 0-1598 µg/g |

Compared to reports of other purple corn varieites, we observed higher percentages of acylated anthocyanins (72%), pelargonidin- (99%) and peonidin-derived anthocyanins (50%), and condensed forms (83%) (Table 1). Total pelargonidin-derived anthocyanins reached nearly 1400 µg/g, cyanidin-derived 872 µg/g, and peonidin 274 µg/g. The lower peonidin content is consistent with our previous findings^1^, but peonidin content is of interest due to its potential to shift hue to a slightly more bluish-red color compared to cyanidin^28^. While cyanidin content is lower than previously reported in other pericarp pigmented purple corn lines, the pelargonidin content is much higher. Currently, orange-red natural colorant sources are limited. Carotenoids can be used in applications requiring orange but are not water soluble and can be expensive and difficult to incorporate into some applications. Carmine, a red pigment derived from cochineal insects, also provides a true red/orange, but can controversial due to its non-vegetarian source. A pelargonidin-dominant, high anthocyanin content purple corn line could provide a much-needed source of water-soluble, acid-stable plant-based red/orange colorant for the food and beverage industry^13^.

Figure 1A illustrates the overall chemical diversity in the AR population using principal component analysis (PCA) based on proportions of total anthocyanin content attributed to each anthocyanin species. An HPLC chromatogram with the peaks used for this analysis can be found in Figure 1B and PCA loadings and peak identities in Table 2. Each anthocyanin was confirmed using retention time and previous mass spectrometry results^14^.

**Table 2:** Summary of variables used for PCA and loadings for PC1 and PC2. Peaks correspond to Figure 2B.

| Peak | Anthocyanin | PC1 | PC2 |
| --- | --- | --- | --- |
| 1 | CaCydG catechin-(4,8)-cyanidin-3,5-diglucoside | 0.144 | 0.404 |
| 2 | CaPgG catechin-(4,8)-pelargonidin-3,5-diglucoside |  | 0.401 |
| 3 | CatPndG catechin-(4,8)-peonidin-3,5-diglucoside | 0.102 | 0.37 |
| 4 | AfCydG afzelechin-(4,8)-cyanidin-3,5-diglucoside | 0.111 | 0.372 |
| 5 | CaCyMdG catechin-(4,8)-cyanidin-3-malonylglucoside-5-glucoside | 0.172 | 0.284 |
| 6 | CaPgMdG catechin-(4,8)-pelargonidin-3-malonylglucoside-5-glucoside |  | 0.158 |
| 7 | AfPgG afzelechin-(4,8)-pelargonidin-3,5-diglucoside | -0.146 | 0.185 |
| 8 | AfPndG afzelechin-(4,8)-peonidin-3,5-diglucoside |  | 0.297 |
| 9 | CyG cyanidin 3-glucoside | 0.35 |  |
| 10 | PgG pelargonidin 3-glucoside | -0.372 |  |
| 11 | AfPgMdG afzelechin-(4,8)-cyanidin-3-malonylglucoside-5-glucoside | 0.269 |  |
| 12 | PnG peonidin 3-glucoside | 0.277 | -0.113 |
| 13 | CyMGs cyanidin malonylglucosides | 0.338 | -0.117 |
| 14 | PgMGs pelargonidin malonylglucosides | -0.331 | -0.145 |
| 15 | PnMG peonidin 3-malonylglucoside | 0.275 | -0.152 |
| 16 | CydiMG cyanidin 3-5-dimalonylglucoside | 0.172 |  |
| 17 | PgMdiGs pelargonidin dimalonylglucosides | -0.321 |  |
| 18 | PndiMG peonidin 3-5-dimalonylglucoside |  |  |
|  | Flavone sum of di-*C*,*C*-hexosyl apigenin, *C*-hexosyl-*C*-pentosyl apigenin, *C*-pentosyl-*C*-hexosyl apigenin, and *C*-hexosyl apigenin | -0.181 | 0.3 |

**Figure 1:**
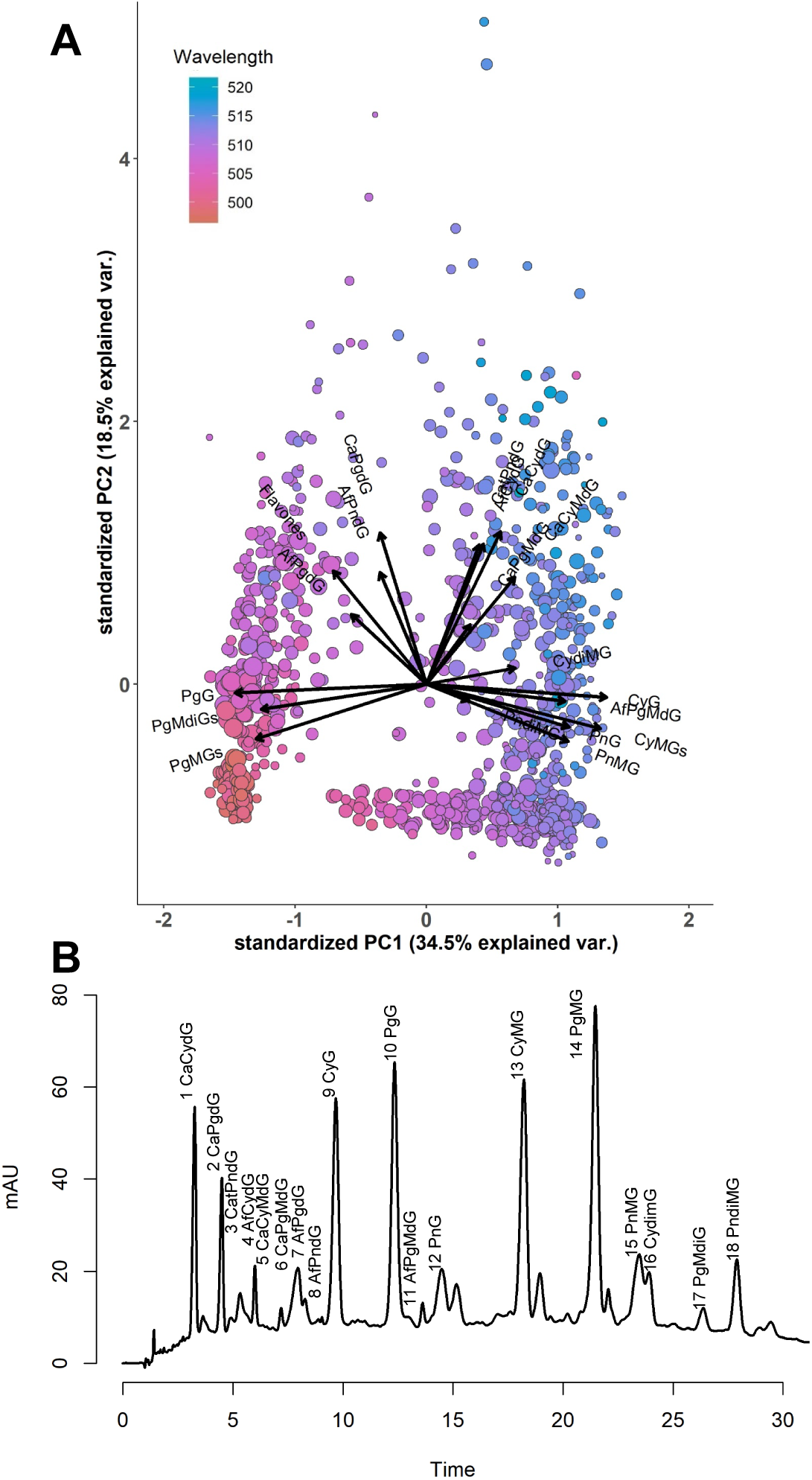
(A) PCA biplot using proportions of total HPLC peak areas and total flavone content. Observations are colored by wavelength (λ_Max_), and size corresponds to total anthocyanin content. (B) HPLC chromatogram showing peaks used for PCA.

The first principal component (PC1) explained 34.5% of the variability in the data set and represents a contrast of pelargonidin versus cyanidin-derived anthocyanins. This is enforced by the color gradient across Figure 1A, in which point color represents wavelength. The λ_Max_ for pelargonidin 3-glucoside (PgG) is ∼496, the λ_Max_ for cyanidin 3-glucoside (CyG) is ∼510^28^, and a gradient from low to high can be seen across the x-axis in the biplot. The second PC explained 18.5% of the phenotypic variability in anthocyanin species and was related to the presence of condensed forms and flavones (high PC2 value) or acylated forms (low PC2 value). Wavelength also appeared to increase with PC2, suggesting condensed forms and flavones may play a role in hue.

### Relationship between anthocyanin profile and λ_Max_

To further analyze the relationship between anthocyanin profiles and hue, λ_Max_ was modeled using PC1 and PC2. PC1, PC2, and their interaction were all significant, which is unsurprising given the color gradients that can be observed in the PCA in Figure 1. We also looked at the major factors associated with PC1 and PC2 based on loadings for each component (Supplementary Figure 3, Table 2). These factors, including proportions of condensed, acylated, and cyanidin/peonidin-derived forms (a measure of pelargonidin versus cyanidin), and flavone content, were plotted against λ_Max_ to identify correlations. These factors were also used to create a complex global model from which the most parsimonious model was selected based on the lowest AIC (Akaike Information Criterion) score, an estimate of model quality. Collinearity between variables and multi-way interactions between factors make interpretation of the model and gauging whether these interactions have a biochemical basis difficult. Nonetheless, cyanidin content especially (p = 1.09e-235), and the other main effects [flavone, p = 7.66e-21; condensed, p = 9.65e-30; acylated (p = 3.67e-8)] were significant, suggesting each can be considered to influence λ_Max_ and overall color in AR lines.

A strong relationship between cyanidin-derived anthocyanins and λ_Max_ was expected, however the correlations with condensed forms and with acylated forms were more surprising. Flavonol-anthocyanin condensed forms have been reported to contribute a bluish hue to wine,^29^ and in purple corn extracts appeared to result in a darker red color but otherwise did not affect color^7^. Acylated and condensed forms are also breeding targets because of their association with anthocyanin stability. Acylated anthocyanins are known to positively impact stability^5^, but reports on the stability of condensed forms are mixed. Some found that lines containing condensed forms produced more stable extracts^6^, while others showed no contribution to stability^7^. Flavone content was also significant in modeling λ_Max_. This is consistent with previous findings^11^ but is our first evidence suggesting that when co-occurring in the same line, C-glycosyl flavone and anthocyanin copigmentation is significant in the determination of extract color. This will likely have major implications on breeding objectives; i.e. when an orange extract is desired, it will be essential to minimize flavone content to avoid an increase in λ_Max_ and a shift from orange to red. Yet when a red-pink or more berry-colored extract is desired, flavone content should be maximized.

While all major anthocyanin factors were significant, interpretation of these results must consider correlations between factors. The proportion of condensed and acylated forms were negatively correlated (r = -0.79, p < 2.2e-16). Some of the condensed forms identified did include acylated groups, and while this could contribute to the correlations, acylated condensed forms were typically present in only very minor amounts. This correlation suggests that acylation and polymerization to make condensed forms are competing processes. Furthermore, if either influenced λ_Max_, the correlation could result in both being significant factors. The proportion of condensed and cyanidin forms were also negatively correlated (r=-0.26, p=5.2e-15). Likewise, this correlation could influence the significance of condensed forms in modeling λ_Max_.

Flavones were positively correlated with condensed forms (r = 0.39, p < 2.2e-16) and negatively correlated with percent cyanidin-derived (r = -0.50, p < 2.2e-16) and acylated anthocyanins (r = -0.29, p < 2.2e-16). While it is possible that these correlations are artifacts of population structure or linkage in the data set, there are also biochemical-based hypotheses that could explain them (Figure 2). For example, it is not unreasonable to hypothesize that the flavonol component of condensed forms, and flavones could share similar regulatory mechanisms, thus explaining their co-occurrence in many of the lines. The relationship between flavones and pelargonidin derived anthocyanins could be explained by substrate specificity. If *Fns1* had a preference for naringenin, a correlation between flavone content and pelargonidin content would be expected, with low flavone concentrations in cyanidin dominant lines. The lack of abundant luteolin-based (eriodictyol-derived) flavones and the exclusivity of apigenin-based (naringenin-derived) flavones supports this hypothesis. Some luteolin compounds were detected with mass spectrometry but were not present in high enough concentrations to adequately measure. More research and intercrossing between AR lines could help determine whether these observed correlations are due to population structure or have biological explanations.

**Figure 2:**
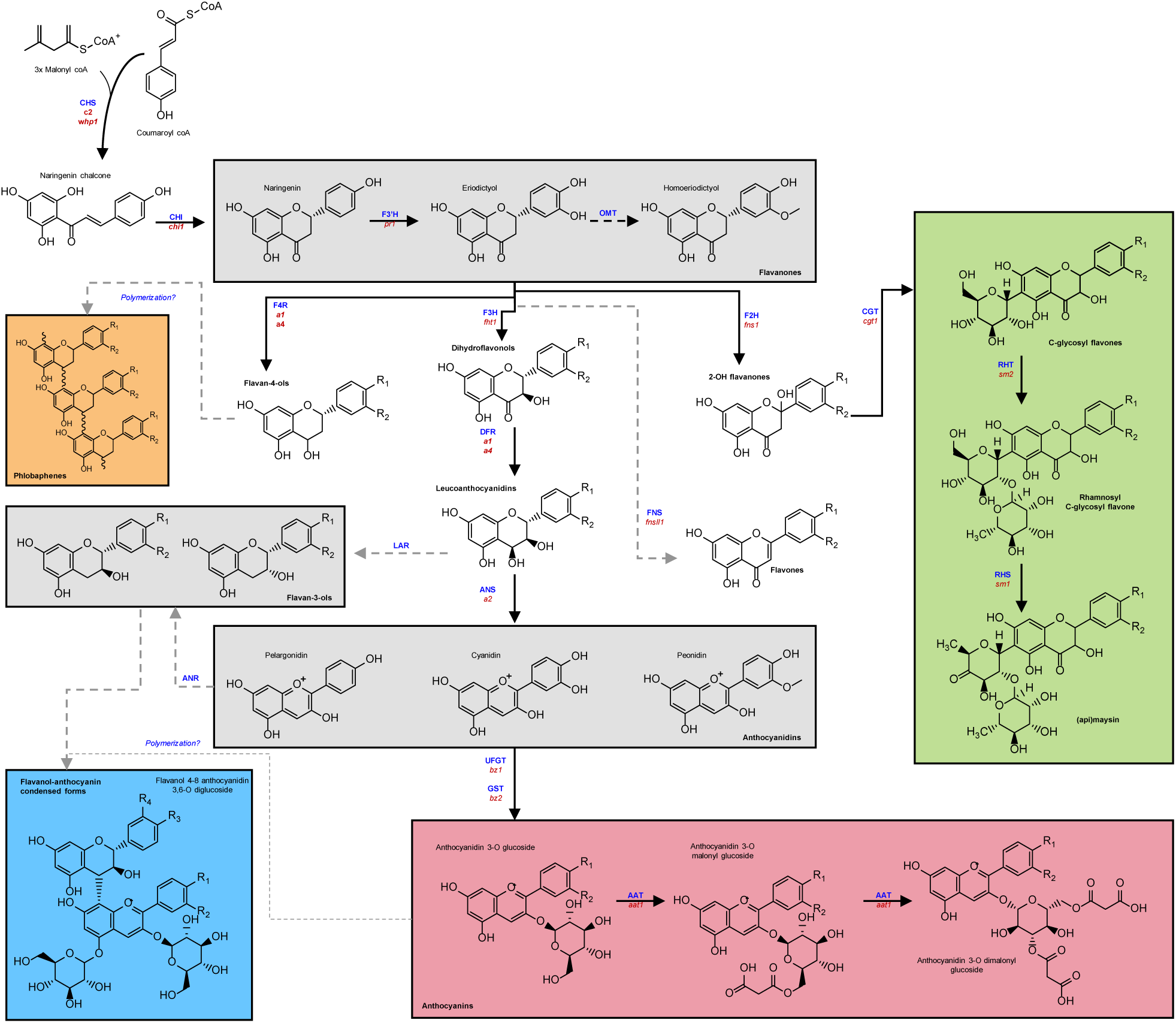
Flavonoid biosynthesis pathway in maize. The pathway stems from phenylpropanoid biosynthesis (upper left). Gray boxes indicate compounds of a similar type that are intermediates in the pathway. Colored boxes represent endpoints of the pathway. Gene product abbreviations (blue) and genes (red) can be found in Table 3.

Analyzing the flavonoid chemistry in Apache Red reveals that extract color is determined by more than just cyanidin to pelargonidin ratio. To successfully breed for specific hues, all of these factors should be considered in a breeding program. However, phenotyping to this extent is time consuming and expensive. Identifying the loci responsible for various anthocyanin decorations and flavones and the ratio between them would be beneficial for eventually creating markers for use in marker assisted selection (MAS).

**Table 3:**
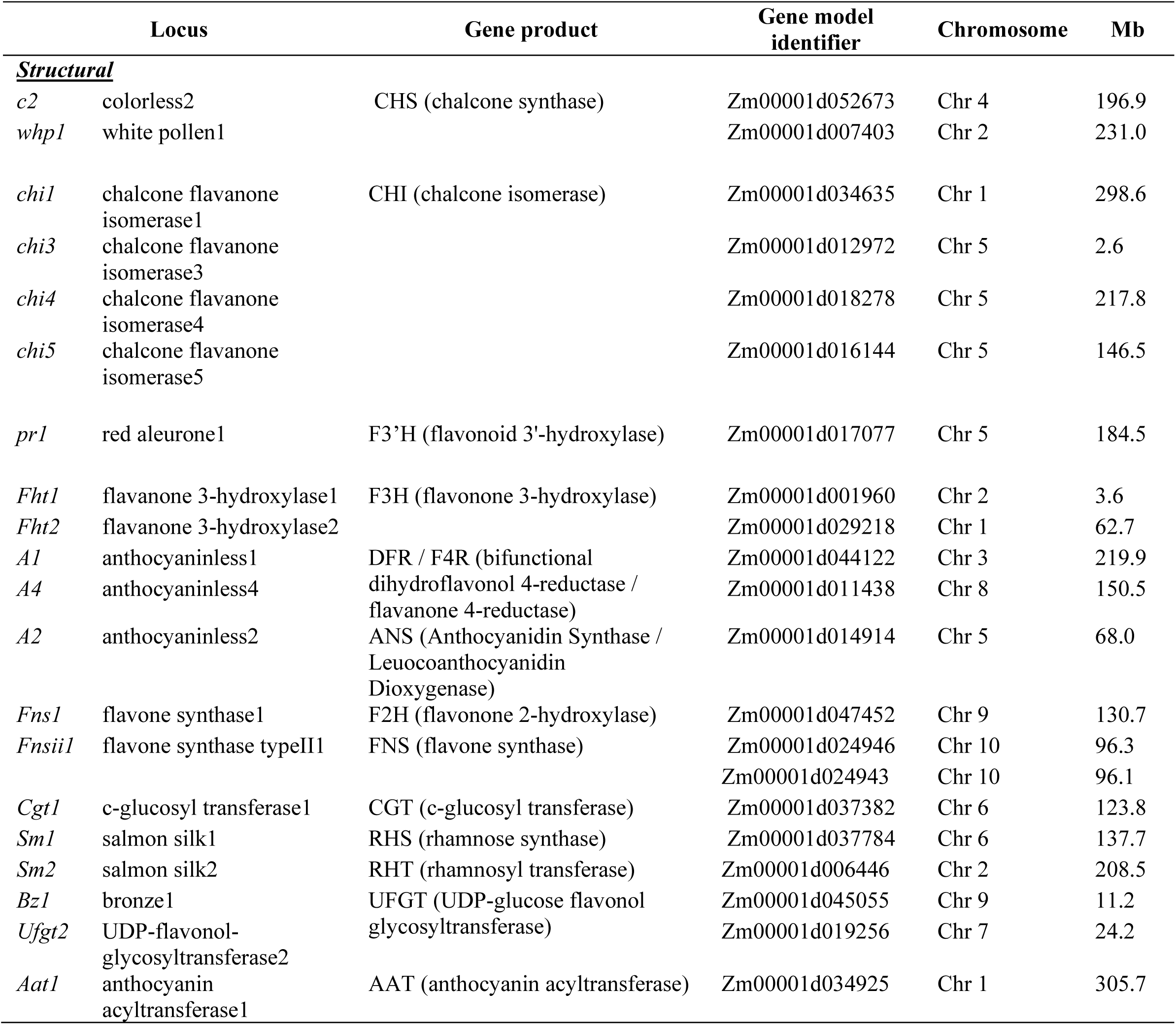
Anthocyanin structural genes corresponding to abbreviated genes and gene products labeled in Figure 2.

### Genotyping and population structure

Given that all the lines were derived from the same landrace, controlling for population structure was a major concern in order to reduce false positives, yet maintainthe power to detect significant associations between genotypes and traits. A PCA of the genotype data for all lines showed that PC1 explained about 10% of the variance in the data set. A drop in the amount of variance explained by each PC (to 2%) was found between PC6 and PC7, and thus 7 PCs were used to help control for population structure in addition to the kinship matrix calculated by GAPIT^21^. To maximize signal detection, we also used a GWAS model that calculates relatedness using a marker subset comprised of all but those on the chromosome where the tested marker resides^30^. Linkage began to decay significantly by 1.5 Mb when calculated by GAPIT, and when a 10 site or 50 site window was used to calculate LD in Tassel, significant decay was seen by 3 Mb and 20 Mb, respectively (Supplementary Figure 4). This is indicative of the number of historical recombination events captured in this study and thus its resolution for marker-trait associations.

### Mapping pelargonidin, cyanidin, and peonidin content

All anthocyanins identified in the AR population were derived from pelargonidin, cyanidin, or peonidin. As discussed above, percentages of these anthocyanidins have a significant effect on extract color, warranting the identification of responsible genetic elements and eventually markers for use in marker assisted selection (MAS). In mapping the proportion of cyanidin-derived anthocyanins, we expected to find *Pr1*, which produces a flavonoid 3’ hydroxylase (F3’H)^9^ in cyanidin containing aleurone lines. However, *Pr1* has not been associated with cyanidin/pelargonidin ratio in pericarp. F3’H catalyzes the conversion of naringenin to eriodictyol and dihydrokaempferol to dihydroquercetin, the steps necessary for producing cyanidin. In maize aleurone, recessive *pr1/pr1* results in the accumulation of almost exclusively pelargonidin-derived anthocyanins and therefore red or pink kernels instead of the purple or blue kernels associated with the presence of cyanidin in most anthocyanin-containing aleurone varieites^8,9,31^. The abundance of pelargonidin-exclusive AR lines and the distribution seen in Supplementary Figure 3D suggests a single gene encoding a F3’H is responsible in pericarp as well.

Indeed, mapping cyanidin-derived anthocyanins resulted in a highly significant peak on Chromosome 5 (Chr 5) near *Pr1* (Figure 3A, Table 4). In aleurone, three nonfunctional *pr1* alleles have been identified, including a 24 nucleotide TA repeat insertion in the 5’ upstream region, a 17 nucleotide deletion near the TATA box, and a Ds insertion in the first exon^9^. Cloning and sequencing of *Pr1* and *pr1* alleles in AR suggests that the same 17 nucleotide deletion observed in aleurone is responsible for nonfunctional *pr1* activity in AR (data not shown). In addition to the highly significant peak near *Pr1*, we also identified several other signals. Most had relatively low minor allele frequencies except for one at the beginning of Chr 8 (<1 Mb). No promising candidates were found for any of these additional loci.

**Table 4:**
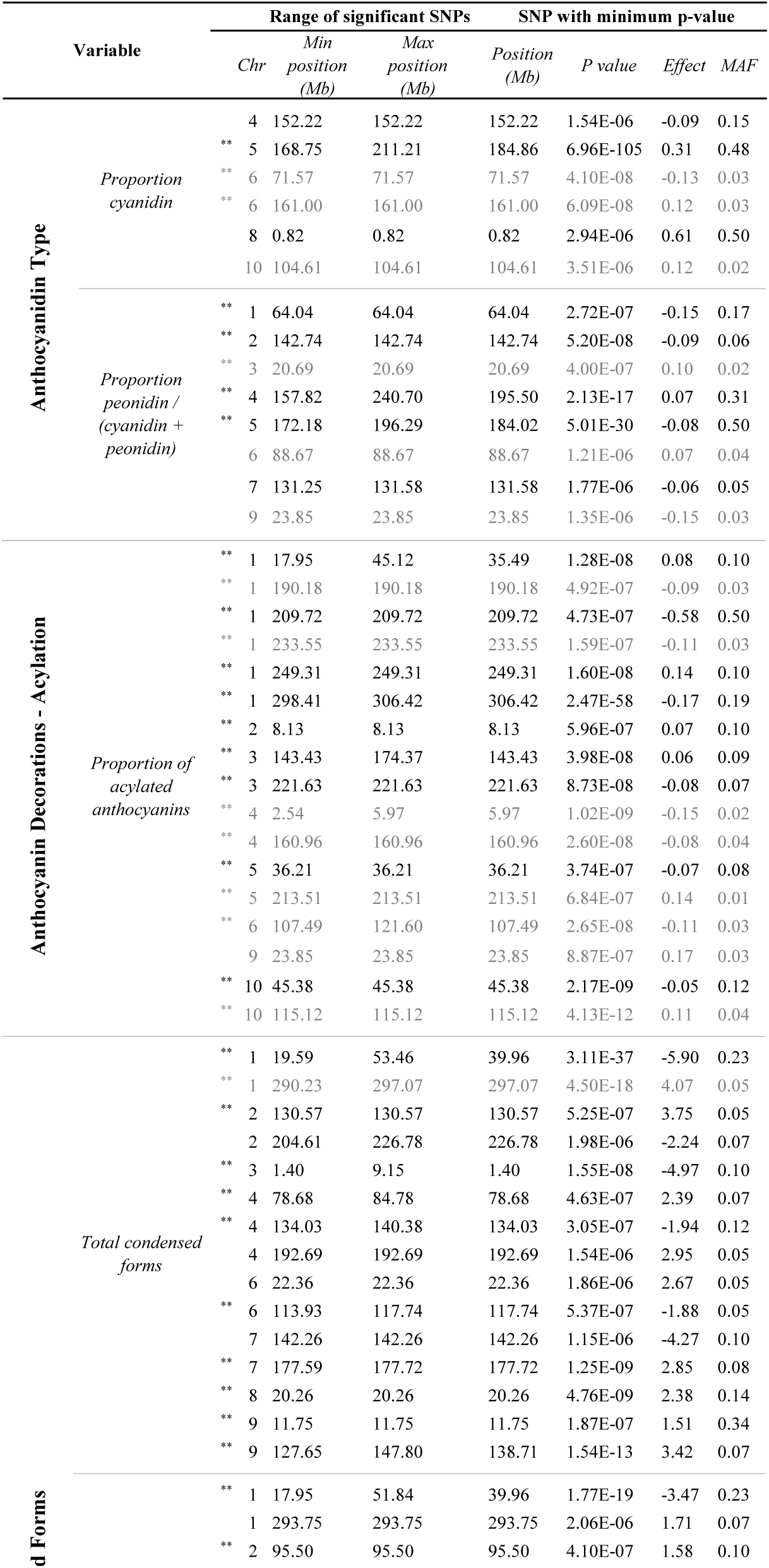
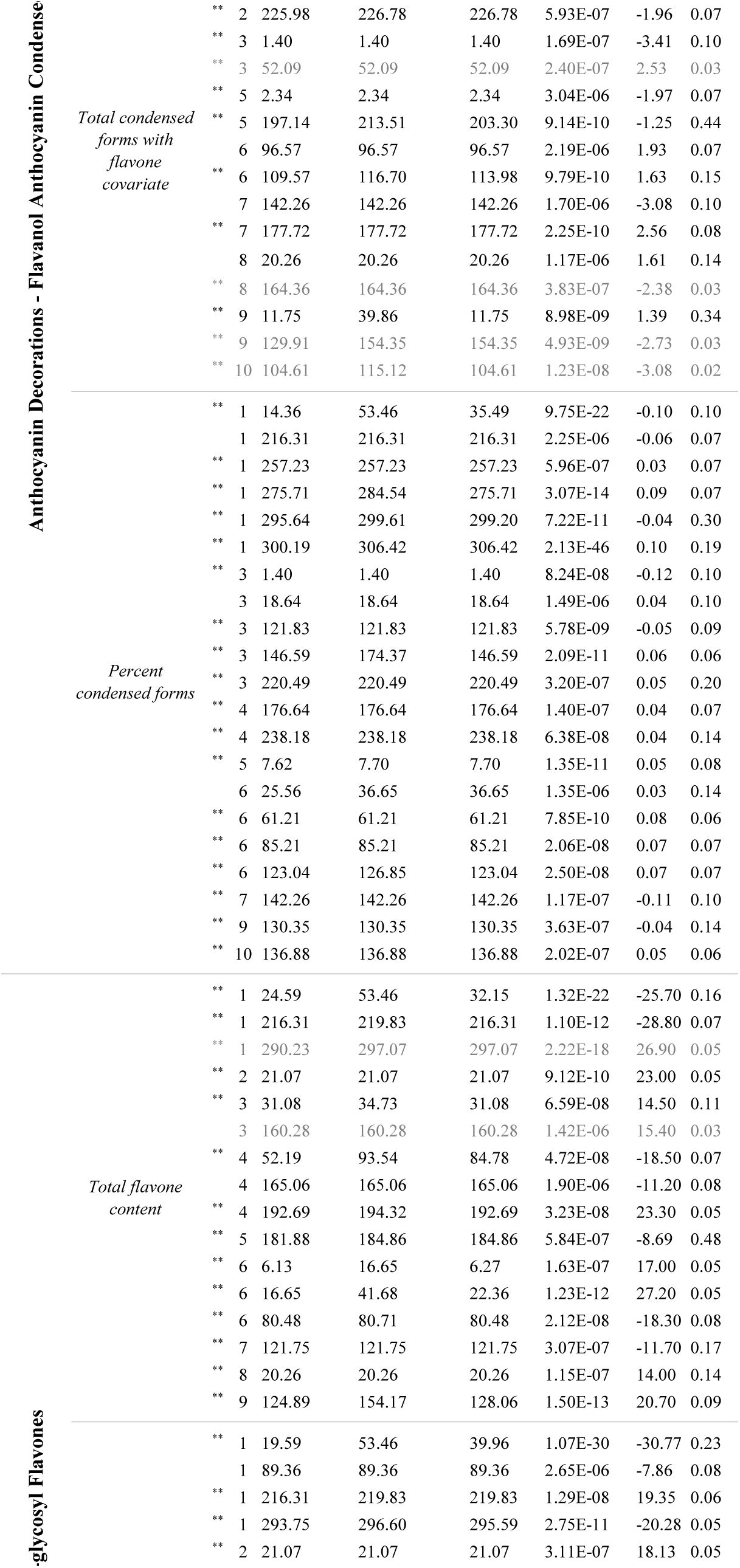
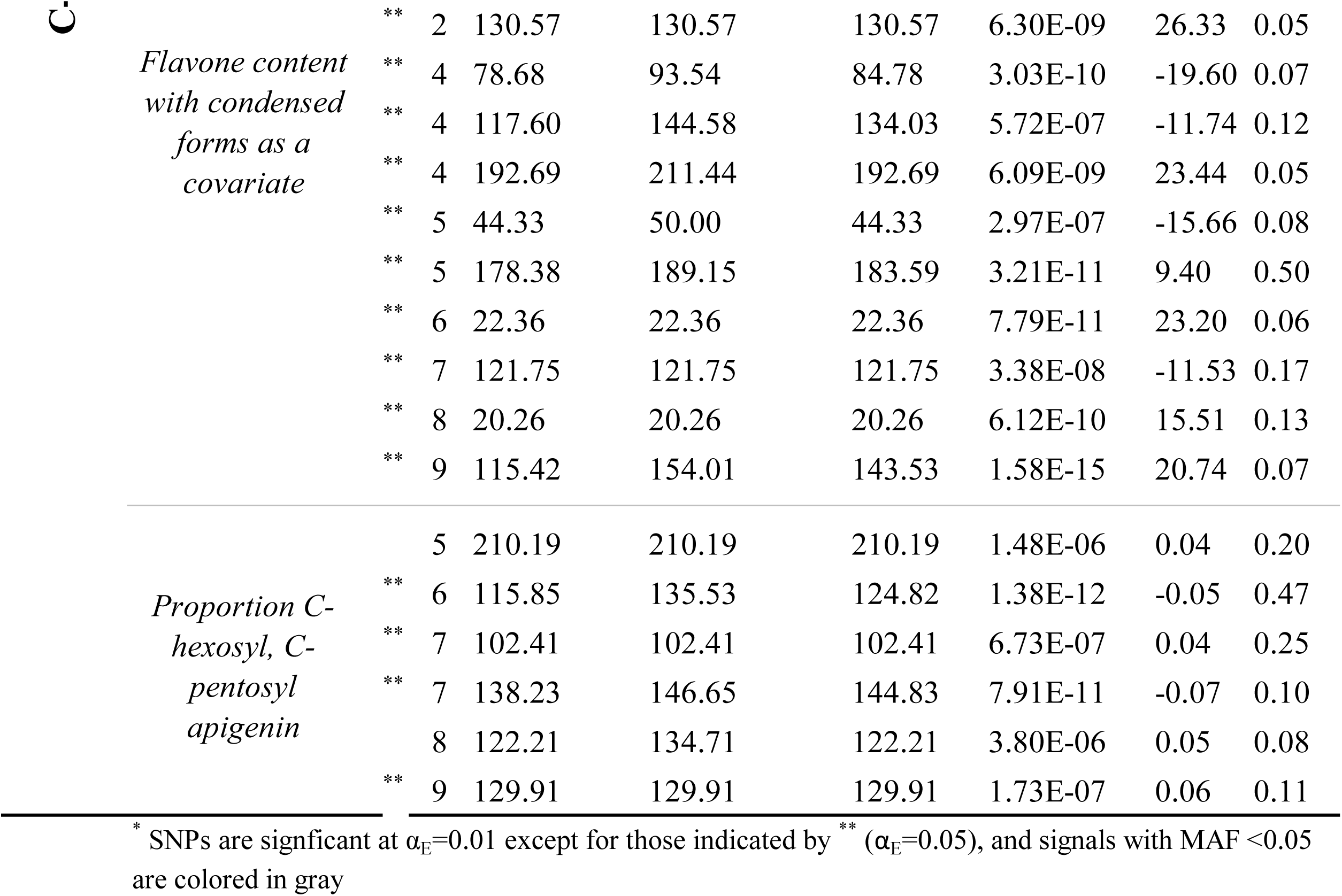
Significant SNPs* for each anthocyanin variable mapped

**Figure 3:**
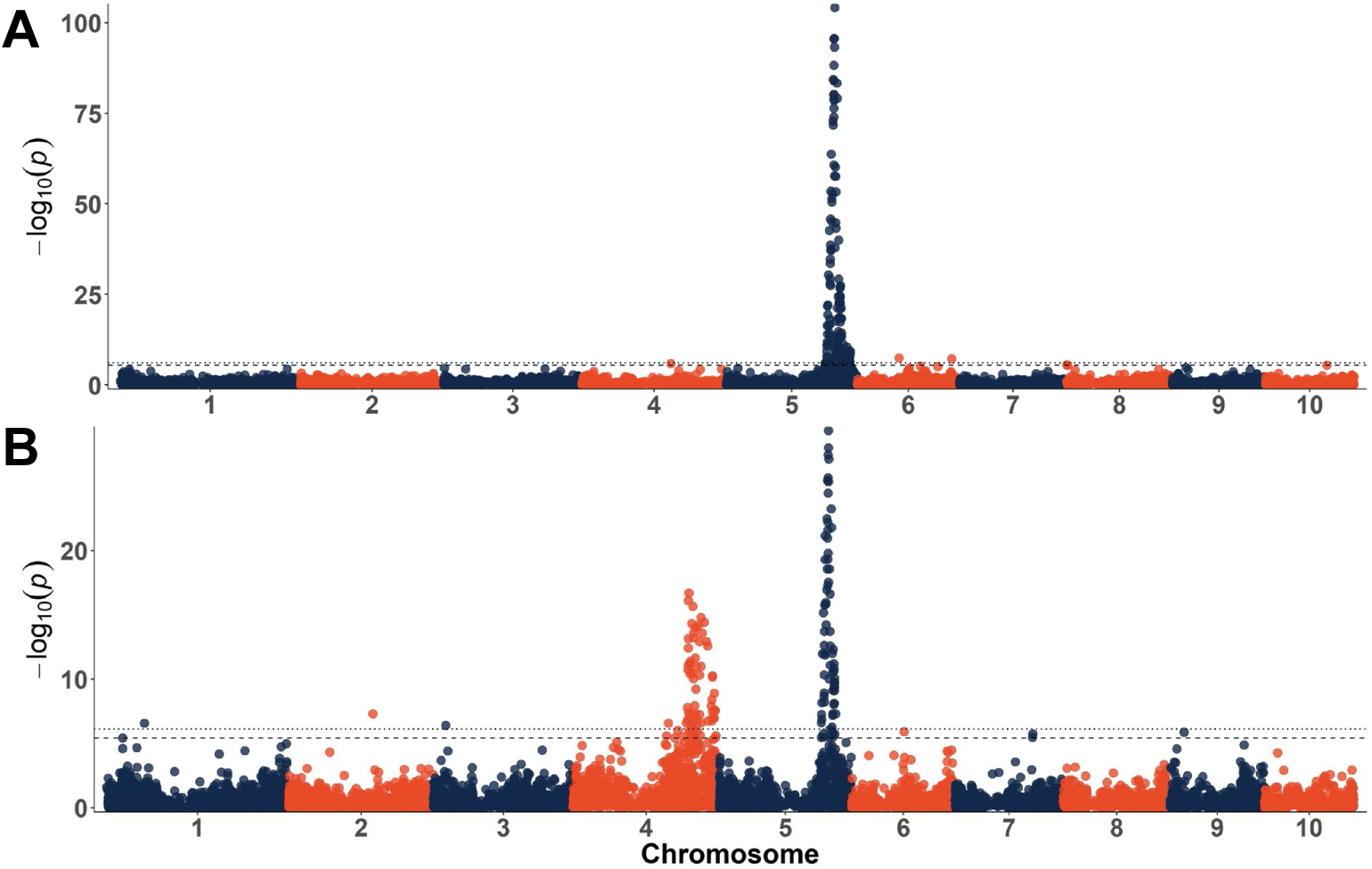
Manhattan plots for percent cyanidin-derived anthocyanins (A) and the percent peonidin of all cyanidin and peonidin-derived anthocyanins (i.e. peonidin / (peonidin + cyanidin)) (B). Dotted lines indicate the experiment-wide significance thresholds using the Bonferroni correction. (α_E_=0.05 and α_E_=0.01)

Mapping the proportion of peonidin-derived anthocyanins produced similar results to mapping the cyanidin proportion - a signal on Chr 5 near *Pr1*(Figure 3B). This is expected since the synthesis of the peonidin precursor, believed to be homoeriodictyol^32^, depends on the conversion of naringenin to eriodictyol catalyzed by the *Pr1* gene product. *Pr1* is likely epistatic with a gene encoding a product with anthocyanin *S*-adenosyl-_L_-methionine-dependent *O*-methyltransferase (AOMT) activity in the formation of homoeriodictyol and thus peonidin.

To isolate AOMT action, we created a new phenotype variable describing the proportion of all eriodictyol-based compounds that were converted to homoeriodictyol-based compounds. This was done by dividing the total peonidin content by the sum of cyanidin and peonidin content. When mapping this variable (Figure 3B), markers near *Pr1* were still significant, but a signal on Chr 4 around 200 Mb emerged. Several other significant SNPs were identified (Table 4), but most had low minor allele frequency, which suggests a higher probability of error. Individual chromosome Manhattan plots showing loci of interest can be found in Supplementary Figure 5.

### Methyltransferase candidates

Many AOMTs have been identified in plants producing O-methoxy anthocyanidins, yet these have all been identified in dicots, and only Caffeoyl CoA *O*-methyltransferase (CcoAOMT) homologs have been identified in monocot species^23^. Recently a CcoAOMT was identified in wheat (*Triticum aestivum*) through transcriptome analysis of purple pericarp. *TaCcoAOMT* was upregulated in peonidin producing wheat pericarp, but the locus was not further characterized^33^. No methyltransferase with AOMT activity responsible for peonidin biosynthesis in maize has been identified, but the presence of peonidin-derived anthocyanins suggests its existence. Amino acid sequences of AOMTs from *Vitis vinifera* and *Glycine max* were blasted against the maize genome^18^, and candidates are listed in Supplementary Table 1. To narrow down the likely candidates, amino acid sequences of AOMTs, CCoAOMTs, and caffeic acid OMTs from other species were aligned with the maize candidates and a maximum likelihood phylogeny was created (Supplementary Figure 6). A clear outgroup containing the caffeic acid OMTs is evident, helping to exclude Zm00001d052683 and Zm0001d052684. Zm00001d052843 and Zm00001d024596 were also in outgroups apart from the remaining AOMTs and CcoAOMTs, which clustered nicely into two groups. Zm00001d045206 and Zm00001d036293 clustered with the CcoAOMTs, but are located on Chr 9 and Chr 6, respectively. The remaining candidates, Zm00001d052841 and Zm00001d052842 were the next most closely related candidates to the AOMTs and CcoAOMTs, are identified as putative CcoAOMTs, and are located on Chr 4 around 202 Mb.

The methyltransferase phylogeny and the proximity to the signal mapped to Chr 4 for peonidin content suggests Zm00001d052841 and Zm00001d052842 are likely candidates. These candidates were aligned to several methyltransferases with known AOMT and CcoAOMT activity, as well as the CcoAOMT expressed in purple wheat pericarp (Supplementary Figure 7). Amino acid residues associated with enzyme function are highlighted and were conserved across all sequences in most cases. Variability was observed between sequences for Arg217 (CoA binding), Tyr219, and Tyr223. Modeling of substrate binding showed that the bulky tyrosine residues present in CcoAOMTs in this location clash with the anthocyanin 3-glucose moiety, i.e. the site cannot accommodate the larger anthocyanin 3-glucose moiety due to obstruction by a bulky tyrosine^23^. In petunia AOMTs, these tyrosines have been replaced by the smaller amino acids, glycine and leucine. In *TaCcoAOMT*, these have been replaced by arginine and leucine, and by isoleucine, leucine and phenylalanine in the two maize candidates. Replacement with phenylalanine certainly would not reduce the size or bulkiness to the same degree that a glycine or leucine substitution would, but a tyrosine to phenylalanine conversion at one of these locations (Tyr219 or Tyr223) was observed for VvAOMT1, VvAOMT3, DhAOMT, and GmAOMT, all of which have anthocyanin OMT activity. Moreover, the relatively low proportions of peonidin (max = 50%) compared to cyanidin (max=86%) observed here and in purple wheat^34^, as well as in other purple corn studies^1^ suggest that the responsible enzyme has relatively low efficiency. If CcoAOMTs were co-opted for anthocyanin use, it is reasonable that they would have suboptimal performance compared to the AOMTs specialized for peonidin production. Here we found higher proportions of peonidin-derived anthocyanins when anthocyanin content was low (r=-0.24, p<2.2e-16), lending support for this hypothesis.

While these residue changes suggest that Zm00001d052841 and Zm00001d052842 could better accommodate the anthocyanin-3-glucoside moiety than the canonical maize CcoAOMTs, it is believed that the methyltransferase in maize acts prior to glycosylation^32^. Evidence for this comes from a study that identified an S-adenosylmethionine-flavonoid 3’-O-methyltransferase capable of utilizing eriodictyol, luteolin, and quercetin as substrates, but not quercetin 3-glucoside in crude protein extracts from maize seedlings, leaf sheath, and aleurone^32^. Comparing Zm00001d052841 and Zm00001d052842 shows that the two are highly similar except for several amino acids and an extra string of 33 amino acids at the beginning of the Zm00001d052841 sequence. Further studies will be required to determine which is responsible for AOMT-like activity, what substrates can be accommodated by the product, and the residues most important for enzyme efficiency.

### Mapping anthocyanin decorations

#### Acylated anthocyanins

Wide ranges of acylated anthocyanins, flavonol anthocyanin condensed forms, and flavones were observed in the phenotypic dataset. Acylated and condensed forms had somewhat bimodal distributions (Supplementary Figure 2), suggesting they are simply inherited traits. Mapping acylated anthocyanin proportions (Figure 4A) produced a peak on Chr 1 near 306 Mb, which aligns with *Aat1*, the anthocyanin acyltransferase recently identified in aleurone pigmented lines and located at 305.7 Mb (Supplementary Figure 5). This helps confirm that *Aat1* is associated with acylated anthocyanin content in pericarp as well as in aleurone. While this was by far the most significant SNP for acylated anthocyanins, several other significant loci were also found and are listed in Table 4.

**Figure 4:**
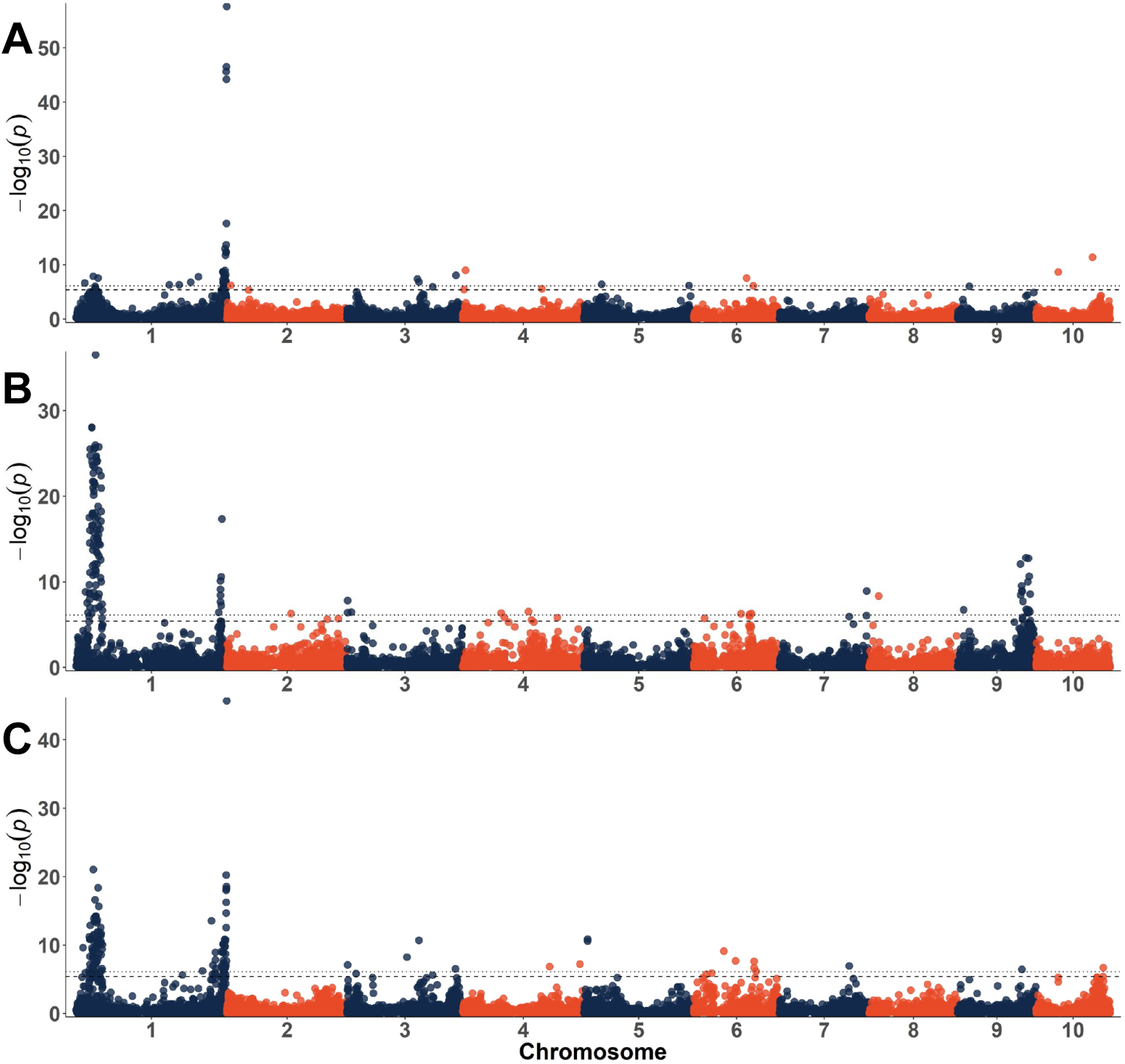
Manhattan plots for anthocyanin decorations: the proportion of acylated anthocyanins relative to the total anthocyanin content (A) total condensed forms (B) and the proportion of condensed forms relative to the total anthocyanin content (C). Dotted lines indicate the experiment-wide significance thresholds using the Bonferroni correction. (α_E_=0.05 and α_E_=0.01)

#### Flavonol-anthocyanin condensed forms

For flavonol-anthocyanin condensed forms, mapping produced a wide signal surrounding *P1*, a known MYB regulator of phlobaphene and flavone biosynthesis^35^ (Figure 4B). We also observed a signal on Chr 9 (127.6-147.8 Mb) near *Fns1* (130.6 Mb), which encodes a flavanone 2-hydroxylase required for flavone biosynthesis. Given the correlation between flavones and condensed forms, finding these loci was not surprising. Several other significant SNPs were also identified (Table 4) but determining whether these correspond to flavone or condensed form biosynthesis is muddied by the correlation between these two phenotypes.

To separate the flavone and condensed form phenotypes, we mapped condensed forms using flavone content as a covariate (Supplementary Figure 8) and looked for both new SNPs and SNPs that remained significant in both analyses. Though drastically reduced in size, the signal on Chr 1 near *P1* was still present. However, the most significant SNP was found at 40 Mb, about 8.5 Mb away from *P1*. The peak on Chr 9 near *Fns1* also remained, but the range of significant SNPs expanded. Along with the phenotypic correlations described, the stability of these signals with the flavone covariate suggests they play an indirect role in the amount of condensed forms accumulated, perhaps by regulating flux through different branches of the flavonoid pathway.

Another signal on Chr 1 was found for both analyses near *Chi1 (Chalcone isomerase1* 298.6 Mb), a flavonoid biosynthesis structural gene shared by both condensed forms and flavones. However the range of the peak changed, with the most significant SNP near 297.1 Mb without the covariate and near 293.8 Mb with the covariate. This moves the signal away from *Chi1* and closer to a potential ANR or LAR (anthocyanidin reductase / leucoanthocyanidin reductase) ortholog (Zm00001d034357) located at 290.5 Mb (Supplementary Figure 9A). Mapping flavonol-anthocyanin condensed forms was anticipated to produce a signal near a candidate LAR or ANR, the enzymes required for the biosynthesis of the flavan-3-ol component of condensed forms (Figure 2). To identify candidates prior to mapping, known LAR and ANR proteins were blasted against the maize genome^18^ and a maximum likelihood phylogeny was created with the hits and orthologs from other species (Supplementary Figure 10). The suggested candidate on Chr 1 is more closely related to the LAR group from this phylogeny. LAR has been associated with the extension of proanthocyanidin polymers of flavan-3-ols^36^, leading to the hypothesis that this same enzyme could be associated with the addition of flavan-3-ols to anthocyanins to make condensed forms.

On Chr 2, significant SNPs were found around 226.78 Mb. The closest candidate we identified is a MYB transcription factor (myb110, Zm00001d007191, 224.25 Mb), which was identified by blasting a transcriptional repressor of anthocyanin and proanthocyanidin genes from *Medicago truncatula* (MtMYB2) against the maize genome^18,37^. A phylogeny containing MYB activators and repressors associated with anthocyanin biosynthesis and potential orthologous maize candidates can be found in the supplementary materials of this article’s companion (Chatham and Juvik, submitted to G3). We also observed a consistent signal at the beginning of Chr 3 near 1.4 Mb, 1.3 Mb from a blast hit for *Arabidopsis TCP3* (*tcptf33*, Zm00001d039371, 2.7 Mb; Supplementary Figure 9E). TCP3 is a transcription factor that interacts with the MYB/bHLH complex to assist in activating flavonoid biosynthesis^38^. A signal near *Pl1* (Chr 6 at 113 Mb), the MYB required for anthocyanin biosynthesis in maize plant tissue, was also found. In addition to the peak on Chr 9 near *Fns1*, we also identified a range of significant SNPs at the beginning of the chromosome (Supplementary Figure 9C). The most significant SNP in this range is less than 1 Mb from *Bz1* (*Bronze1*) a UDP-glucose flavonol glycosyltransferase responsible for glycosylating anthocyanins^39^. Given the wide range created when the flavone covariate was added, other candidates may include a flavonoid 3-*O-*glucosyltransferase (Zm00001d045254) and *ZmMrp3*, an ABC transporter associated with transport of anthocyanins to the vacuole. However both of these are over 4 Mb away from the most significant SNP. This signal could also be tagging *Bp1* (*Brown pericarp1 -* located in bin 9.02 (11.8 – 23.3 Mb in B73 RefGen_v3)), a locus that has been suggested to play a role in the polymerization of flavan-4-ols to create phlobaphenes and is epistatic with *P1*^12^. No gene model has been associated with the *Bp1* locus, but if polymerizing activity is confirmed, it could participate in the heterodimerization between flavan-3-ols and anthocyanins to make condensed forms. Conversely, its potential regulation of flavone biosynthesis could explain why it was found for condensed forms since flavones and condensed forms are correlated in this population. Other signals observed consistently between the analysis with and without the flavone covariate were found on Chr 7 around 142 Mb and 178 Mb, each with relatively large effect sizes. A potential candidate was found at 138.2 Mb (LAR/ANR ortholog - Zm00001d020970), but this is over 3 Mb away and the signal observed was comparatively weak. A glycosyltransferase annotated as an anthocyanidin 3-O-glucosyltransferase (Zm00001d022467 at 178.22 Mb) was found close to the signal at 178. Other significant SNPs detected can be found in Table 4.

Several new signals were identified when the flavone covariate was used. A peak on Chr 5 was found near 203.3 Mb, 1.5 Mb from *Pac1* (*Pale aleurone color1*) and 3 Mb from an ANR/LAR candidate (Zm00001d017771; Supplementary Figure 9D). *Pac1* is a WD40 repeat protein and member of the ternary MBW complex that regulates anthocyanin biosynthesis^40,41^, but *Pac1* is only required for complete anthocyanin production in aleurone, not vegetative tissue^42^. Whether *Pac1* functions in pericarp has not yet been determined. The other candidate, Zm00001d017771, groups with orthologous LARs (Supplementary Figure 10). New signals on Chr 8 (164.4 Mb) and Chr 10 (104.6 Mb) were also found but no potential candidates were identified, and both had minor allele frequencies below 0.05.

In addition to these analyses we also mapped condensed forms as a proportion of the total anthocyanin content (Figure 4C). Several signals were redundant with mapping total condensed forms, including signals near *P1, Chi1, tcptf33, Fns1*, and at Chr7:142.26 Mb. *Aat1* was also previously identified for the proportion of acylated anthocyanins but given the inverse relationship between the proportion of acylated and condensed forms, its recurrence here is not surprising.

Though relatively weak, a signal on Chr 1 was found less than 1 Mb away from *myb83*, a transcription factor with similarity to *AtMYB60*, which functions as a repressor of anthocyanin biosynthesis^43^. We also found peaks on Chr 1 and Chr 5, both near UDP-D-glucose dehydrogenases with potential roles in sugar metabolism and expression in pericarp tissue^35^. On Chr 3, a peak near the *A1* (*Anthocyaninless1)* locus was identified. *A1* encodes a dihydroflavonol 4-reductase required for anthocyanin biosynthesis. Another candidate for this signal may be *mybr97*, a protein with similarity to MYB repressors from Petunia (*MYBx*) and *Arabidopsis* (*CPC*). CAPRICE (CPC) is an R3-MYB repressor that negatively regulates anthocyanin biosynthesis by competing for bHLH binding^44^. Similar repressors were identified in petunia (*MYBx)* and tomato (*Atroviolacea* (*Atv*))^45^. *Mybr97*, located at 221.6 Mb on Chr 3 is close to the *A3* locus (between 205.4 Mb and 215.9 Mb based on B73_refgen_V2) identified as a negative regulator of anthocyanin biosynthesis and for which no gene models have been associated.^46^ On Chr 4, significant SNPs were found at 176.6 and 238.2. The best candidates identified for these QTN were a blast hit for an ATPase from petunia (*PH5*) involved in flower color^47^ and a vacuolar proton pump (vpp3) (Zm00001d052022 and Zm00001d053765, respectively). *PH5* influences the vacuolar pH and thus flower color and is required for the accumulation of proanthocyanidins. Since both proanthocyanidins and condensed forms require flavan-3-ols as building blocks, it is plausible that an ATPase in maize could be required for effective transport of flavonols and thus the synthesis of condensed forms^41^.

On Chr 6 we identified several significant SNPs that appear to be on either side of the centromeric region. Stronger LD in this region likely limits our ability to locate with much certainty the region containing a causative SNP. Nonetheless, we found a potential MYB candidate close to one of the QTNs. This candidate, *myb1* (Zm00001d035918) was a close match when *CPC* and *AtMYB60*, both repressors of anthocyanin biosynthesis in *Arabidopsis*, were blasted against the maize genome^18,44^. Additionally, a signal on Chr 10 was found about 3 Mb away from *R1*, the bHLH component of the MBW ternary regulatory complex typically associated with activating anthocyanin biosynthesis in aleurone^41^, Originally, mapping condensed forms was expected to produce several large effect loci, especially given the current understanding of the pathway (Figure 2) and the somewhat bimodal distribution observed for condensed forms (Supplementary Figure 3b). However, the regulation of condensed forms appears to be a rather complex, quantitative trait, at least in this AR population.

#### Mapping flavone content

C-glycosyl flavones can largely influence anthocyanin extract hue and intensity and are therefore an important consideration in a breeding program aiming to optimize hue^11^. They are also the precursors to maysin, a C-glycosyl flavone that provides resistance to corn earworm (*Helicoverpa zea*)^48^. Surprisingly, we were not able to identify significant quantities of maysin in the AR population, suggesting fixation of a null *Sm1* (*Salmon silks1)* or *Sm2* (*Salmon silks2)* allele. *Sm1* and *Sm2* encode the rhamnose synthase and rhamnose transferase that convert *C*-hexosyl flavones into maysin or apimaysin^49^ (Figure 2). As determined by HPLC and confirmed using mass spectrometry^14^ the major C-glycosyl flavones present in AR lines were di-*C, C*-hexosyl apigenin, *C*-hexosyl-*C*-pentosyl apigenin, *C*-pentosyl-*C*-hexosyl apigenin, and *C*-hexosyl apigenin.

Mapping flavone content was performed with and without a covariate for condensed forms, but many of the same loci were detected as for condensed forms (Table 4). As expected, a large signal near *P1* was identified(Figure 5A). However, the signal was wide, ranging from 20 Mb to 53 Mb with the most significant SNP around 40 Mb, more than 8 Mb away from *P1*. Several loci involved in sugar metabolism are near *P1* and are regulated by it^35^. While it’s possible that this signal represents multiple signals, it is equally possible that linkage blocks in this area are contributing to the signal width (Supplementary Figure 11). In addition to this major locus, a signal was identified on Chr 1, 1.5 Mb from *Chi1*, which encodes a chalcone isomerase that functions prior to the branch point in the pathway for flavones or anthocyanins. We also detected a peak around 216 Mb found previously when mapping the proportion of condensed forms (Supplementary Figure 12).

**Figure 5:**
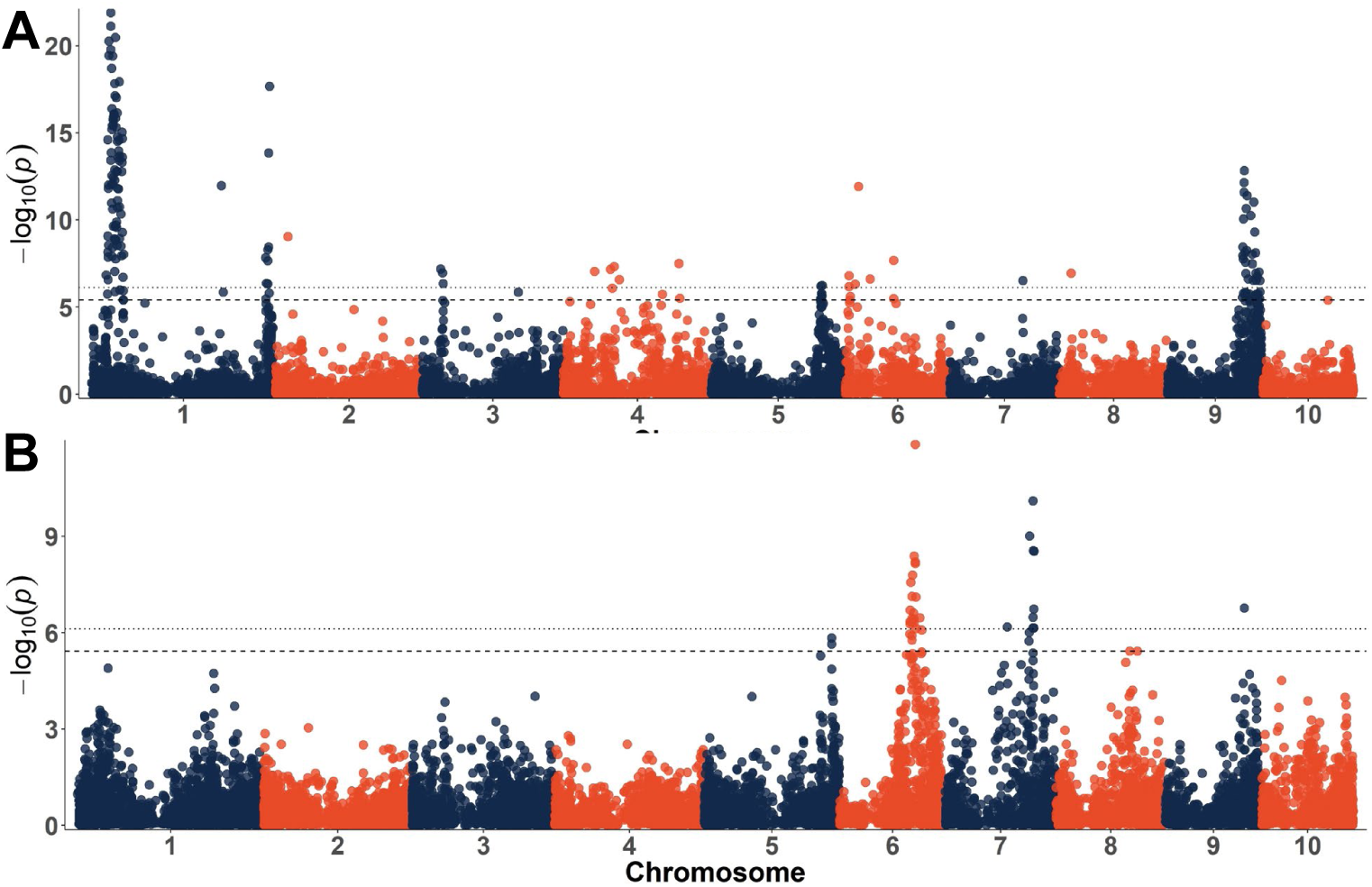
Manhattan plots for total C-glycosyl flavones (A) and proportions of C-hexosyl,-C-pentosyl apigenin (B). Dotted lines indicate the experiment-wide significance thresholds using the Bonferroni correction. (α_E_=0.05 and α_E_=0.01)

While not detected for condensed forms, we also found an isoflavone reductase-like protein on Chr 3 (*Irl1*, Zm00001d040173, 30.6 Mb) which is a close blast match to both the LAR from *Medicago* and an LAR from *Desmodium uncinatum* (a member of the isoflavone reductase-like group of the reductase-epimerase-dehydrogenase protein family) (Supplementary Figure 9)^50,51^. While this candidate may be more appropriate for the biosynthesis of condensed forms than flavones as a response variable, this locus should not be overlooked as a strong candidate for a maize orthologous LAR due to the correlation between flavones and condensed forms in this population. Also unexpected was a relatively weak signal on Chr 2, 2 Mb away from *B1*, a plant-specific MYB activator of anthocyanin biosynthesis^52^. If this locus contributed to flux toward the anthocyanin side of the pathway, it could influence flavone content as well given these two compound classes share common precursors.

Unsurprisingly, we also found a strong signal near *Fns1*, the flavone synthase required for flavone biosynthesis in maize^53^. This confirms both that the same pathway is functioning in Apache Red, as well as the power of this dataset to detect loci underlying flavonoid biosynthesis despite the number of strong correlations between response variables. Also expected, was a signal near *Pr1.* The majority of flavones detected in AR were apigenin-derived, containing a single hydroxyl group on the B-ring. Rather than an inability to produce luteolin compounds (two hydroxyl groups on the B-ring) though, we suspect this is more likely due to trait linkage resulting from a fixed recessive *pr1* in AR families that produced large quantities of flavones. Intercrossing between *Pr1/-* lines (evident by cyandin-dominant anthocyanins) and high flavone lines would likely produce luteolin-derived flavones; however, this should be tested to confirm. Production of different flavone types in AR may also be useful for investigating the copigmentation properties of various flavone-anthocyanin combinations. Other signals identified when mapping flavones, but without promising candidates, are listed in Table 4.

With data quantifying different flavone species from HPLC analysis we were also able to map the proportion of each flavone type relative to total flavone content. We again identified a signal near *Fns1*, as well as one on Chr 6 near *Cgt1*, a dual *C-/O-*glycosyltransferase implicated in the formation of *C-*hexosyl flavones^54^. *Cgt1* is capable of *C-*glycosylating at both the 6-*C* and 8-*C* positions, but this is the first indication of its role in creating di-*C,C-*hexosyl flavones, glycosylated at both positions on the same molecule. While it is possible that *Cgt1* is also capable of *C-*glycosylating pentose moieties, it is difficult to separate the functions in this data set as no *C-*pentosyl flavones were identified that did not also contain a hexose. In addition to these signals, peaks were found on Chrs 5, 7, and 8, which are listed in Table 4. No obvious candidates were identified for these, but several plausible loci were considered. For the peak on Chr 7 with the most significant SNP located at 144.83 Mb and spanning 138–146 Mb, two glycosyltransferases were found around 144 Mb (Zm00001d021167 and Zm00001d021168). This signal is also close to *myb152* (148.15 Mb) a positive regulator of the phenylpropanoid pathway^55^. We also found a MYB transcription factor close to the signal on Chr 8 (*myb147*) as well as a flavonoid 3’-hydroxylase orthologous to Transparent Testa 7 (*TT7*), which functions similarly to *Pr1* in maize, determining the ratio of kaempferol to quercetin in Arabidopsis seed coats^56^. A table and phylogram of candidate MYBs and known flavonoid MYBs from other species can be found in the supplementary materials of Chatham and Juvik, submitted to G3. Variations in flavone structure could influence copigmentation efficacy^27^, however more information is needed before specific breeding objectives for optimal flavone content can be defined.

#### Applications

The development of separate flavone-rich lines and anthocyanin-rich lines is warranted to allow for the maximum range of applications. Ideally, a pelargonidin-rich line producing orange extract, a cyanidin-rich line producing red-pink extract, and a colorless line rich in C-glycosyl flavones with the greatest capacity for copigmentation would be bred. Doing this would allow a manufacturer to design a range of hues for use in food and beverage products by mixing different ratios of the pelargonidin extract, cyanidin extract, and flavone extract, as described previously^11^. Anthocyanin-rich lines should be further optimized to maximize stability and anthocyanin content. Acylated anthocyanins should be maximized, while condensed forms should be minimized. Reports of improved stability from condensed forms are sparse, and condensed forms were associated with flavone content in this population, something that should be minimized in breeding anthocyanin lines. Condensed forms could also represent a reduction in overall content, since two units (1 flavonol + 1 anthocyanin) are required to make one condensed form.

The development of markers for the candidates and known genes identified herein to be associated with anthocyanin type and decoration could improve selection efficiency. For example, pelargonidin-rich lines could be selected at the seedling stage by choosing *pr1/pr1* lines and could be tracked through backcrossing by differentiating between *Pr1/pr1* and *Pr1/Pr1* individuals to avoid subsequent test crosses ensuring maintenance of the recessive allele). Combined with markers for *Pr1*, the identification of a methyltransferase candidate for peonidin content could enable the development of markers that aid in creating peonidin-dominant lines (requiring both *Pr1/-* and a functional *AOMT* due to their complimentary gene action). Based on results presented here, purple corn breeding programs should select for nonfunctional *p1* and nonfunctional ANR or LAR candidates to eliminate flavones and condensed forms while selecting for functional *Aat1* to maintain acylated anthocyanins.

#### Conclusions

Together, the diverse range of flavonoids present in AR provide an assortment of specialized metabolite profiles that can be modified through breeding. Variation in flavonoid profiles may lead to wider color ranges, improved stability, and intensified color for the use of purple corn in foods and beverages.

This survey provides a comprehensive analysis of anthocyanin species diversity in purple corn and highlights the importance of AR as breeding material in programs selecting for natural colorants. Through mapping, we have provided evidence to confirm the function of *Pr1* and *Aat1* in pericarp in addition to aleurone, a fact that has been assumed but not previously tested. Furthermore, several candidate loci involved in the partitioning of various anthocyanin species affecting the resulting extract hue and stability were identified. Among these, a candidate methyltransferase was identified and believed to be associated with peonidin content. If confirmed with additional studies, this would be the first characterized methyltransferase with anthocyanin *O-*methyltransferase-like activity in a cereal. This study also represents the first investigation of genetic factors underlying the formation of flavanol-anthocyanin condensed forms. The candidate genes identified here will provide a foundation for future research on loci associated with anthocyanin species and decorations.

## List of Abbreviations Used

ABC: ATP-Binding Cassette,
AR: Apache Red,
bHLH: basic Helix-Loop-Helix,
GBS: Genotyping-By-Sequencing,
HPLC: High-Performance Liquid Chromatography,
MAF: Minor Allele Frequency,
MATE: Multidrug and toxic compound extrusion
MYB: derived from “Myeloblastosis protein”,
PCA: Principal Component Analysis,
SNP: Single Nucleotide Polymorphism,
QTL: Quantitative Trait Loci.

## Declarations

### Ethics Approval

Not applicable

### Consent for Publication

Not applicable

### Availability of Data and Material

Lines from the Apache Red population are available by request. All genotype and phenotype datasets will be made available at the time of publication via Figshare under project name “Linking anthocyanin diversity, hue, and genetics in purple corn”.

### Author’s Contributions

LAC developed, genotyped, and phenotyped the Apache Red population, analyzed the data, and prepared the manuscript. JAJ assisted in conceiving the study and revising the manuscript. All authors read and approved the manuscript.

## Acknowledgements

We would like to thank Dr. Pat Brown for assisting with GBS library construction.

## Tables and Figures

**Supplementary Table 1:**
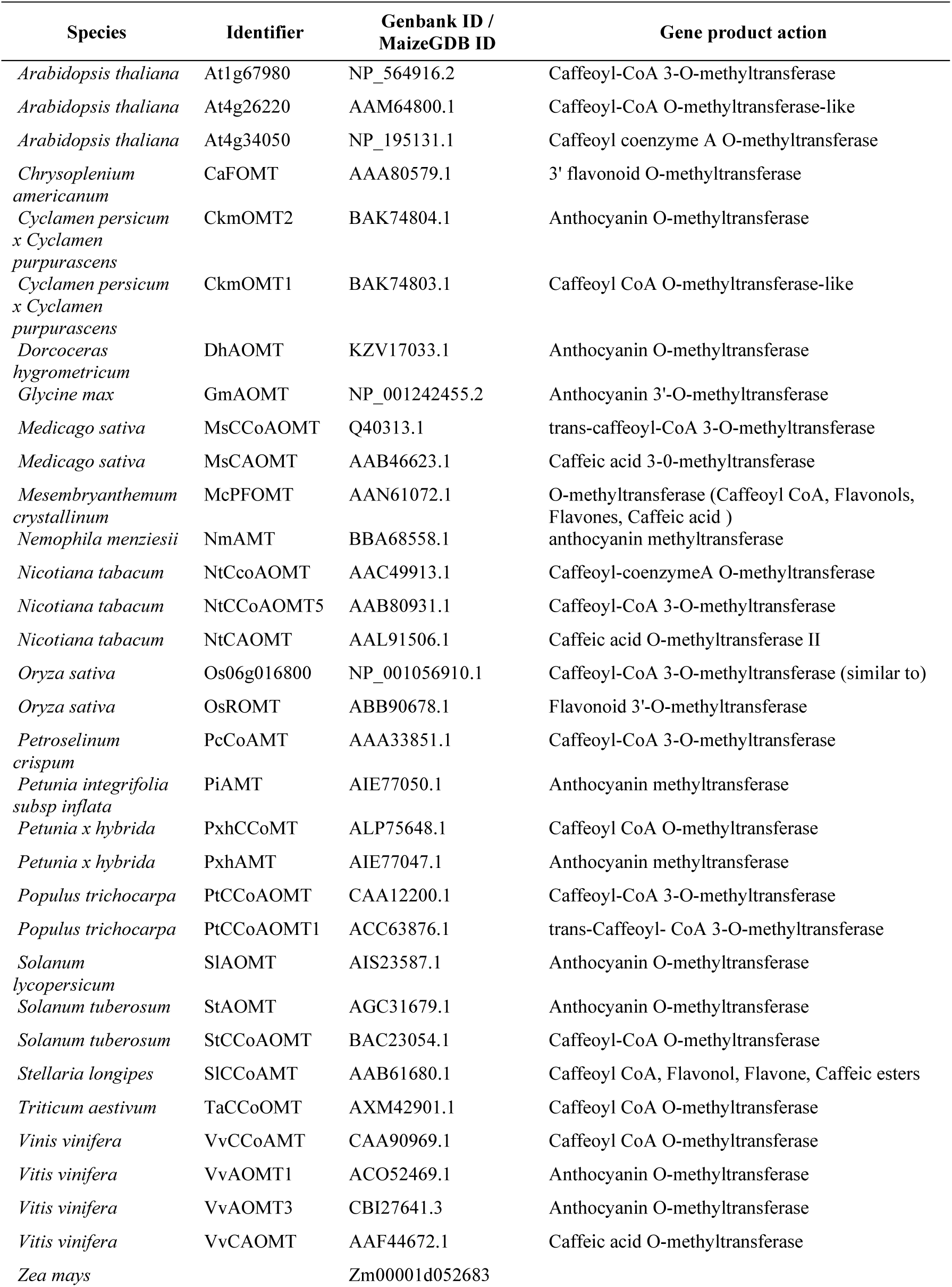
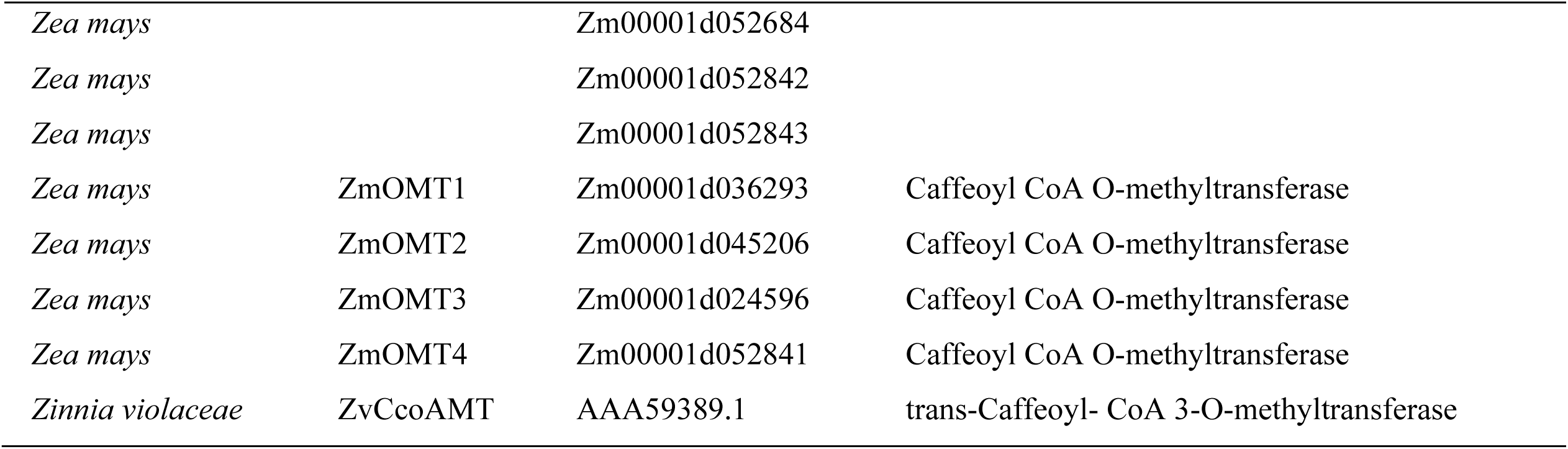
Description of proteins used for methyltransferase phylogram (Supplementary Figure 6) and alignment (Supplementary Figure 7).

**Supplementary Figure 1:**
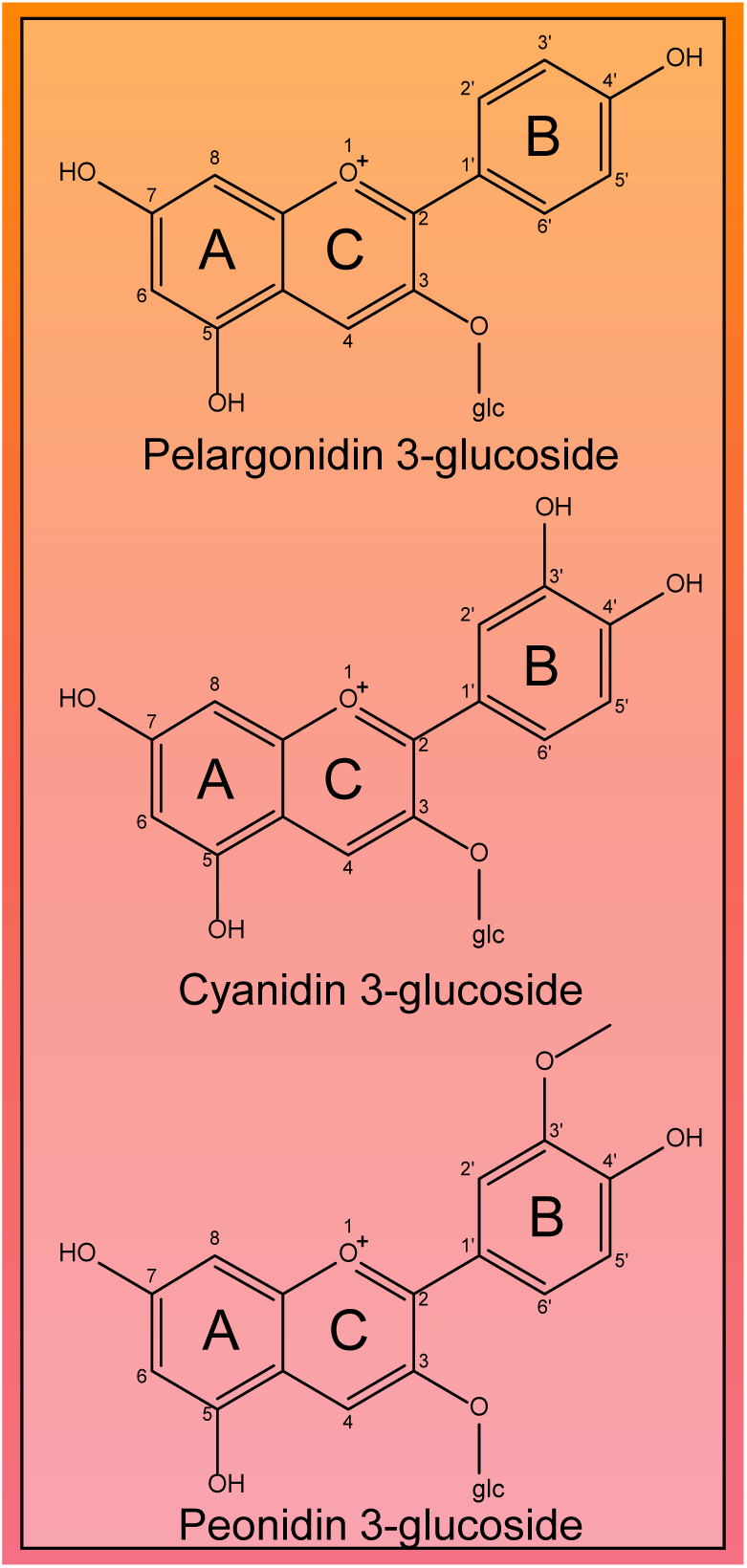
Anthocyanin glycoside structures.

**Supplementary Figure 2:**
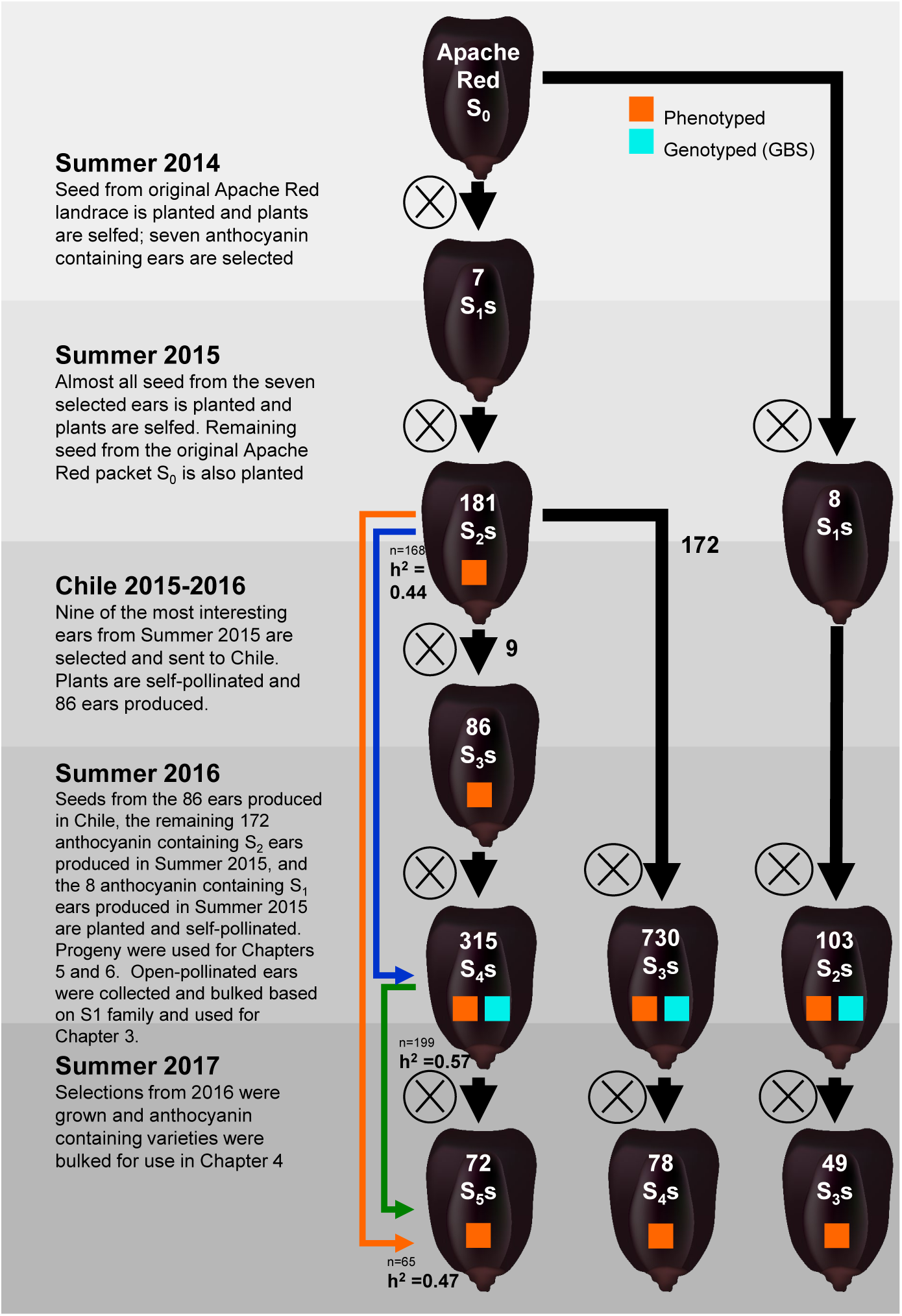
Schematic for the creation of Apache Red lines.

**Supplementary Figure 3:**
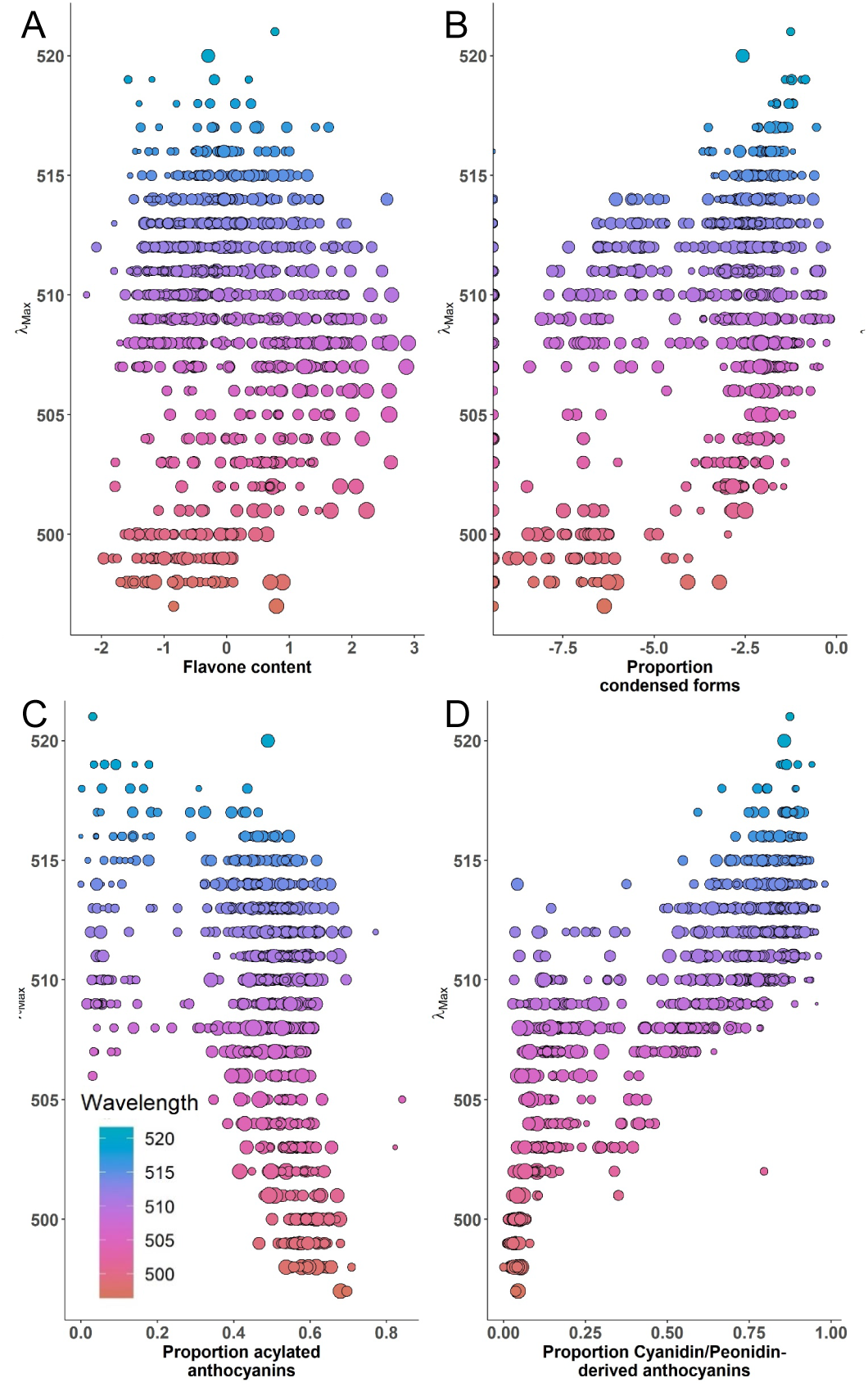
Correlations between anthocyanin composition factors and wavelength at maximum absorbance (λ_Max_), a proxy for hue.

**Supplementary Figure 4:**
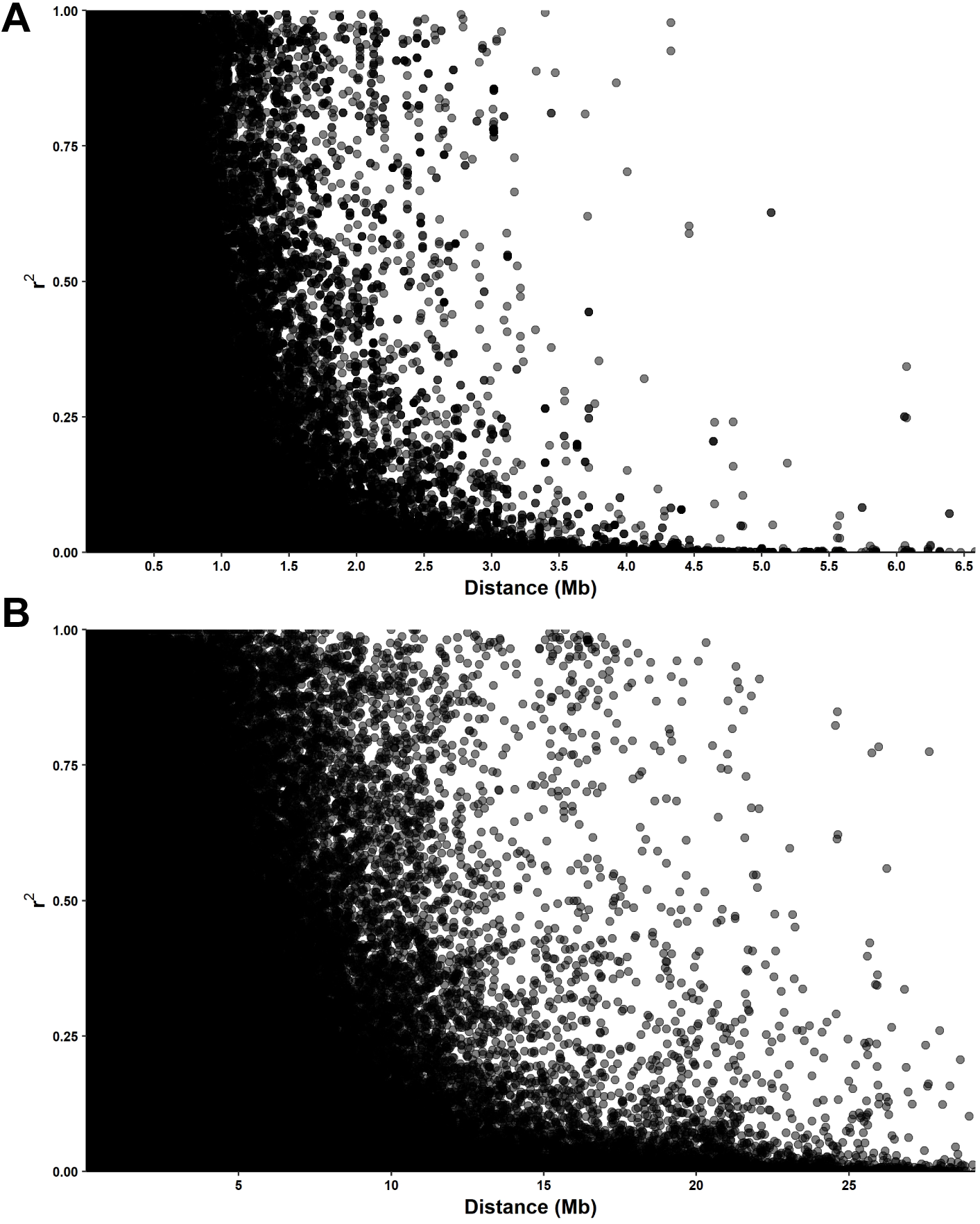
LD decay using r^2^ with either a 10 site window (A) or 50 site window (B)

**Supplementary Figure 5:**
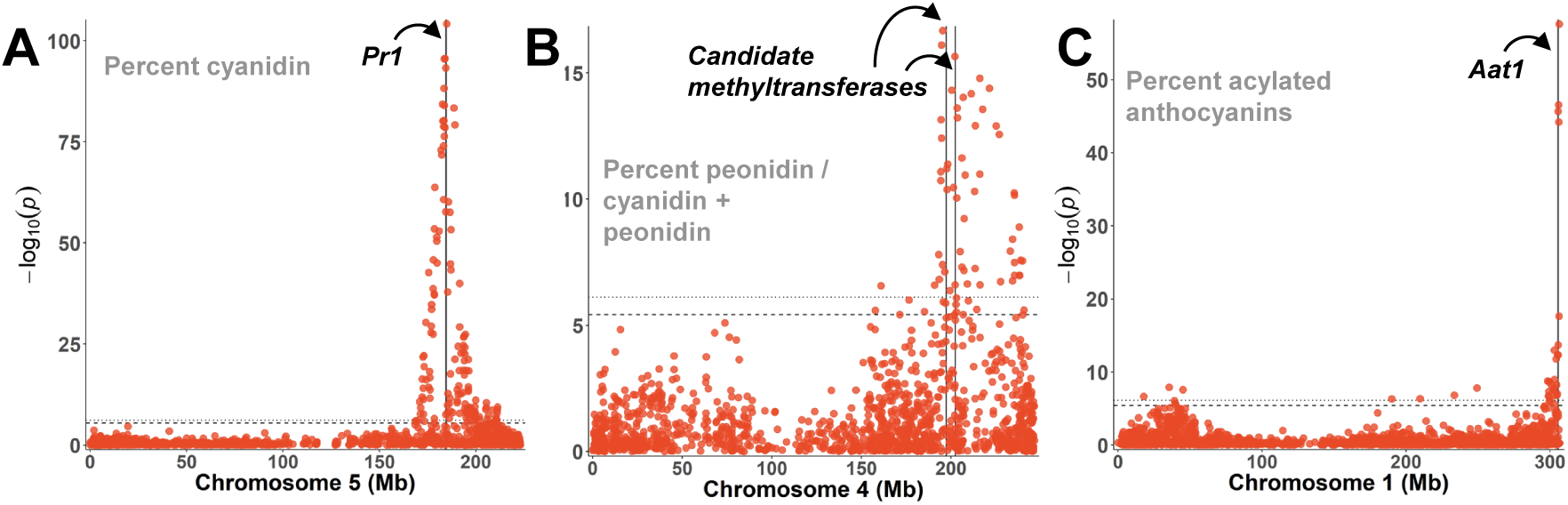
Individual chromosome plots for major signals identified for the proportion of cyanidin (A), peonidin (B), and acylated anthocyanins (C). Chromosome numbers are listed in the x-axis label for each plot.

**Supplementary Figure 6:**
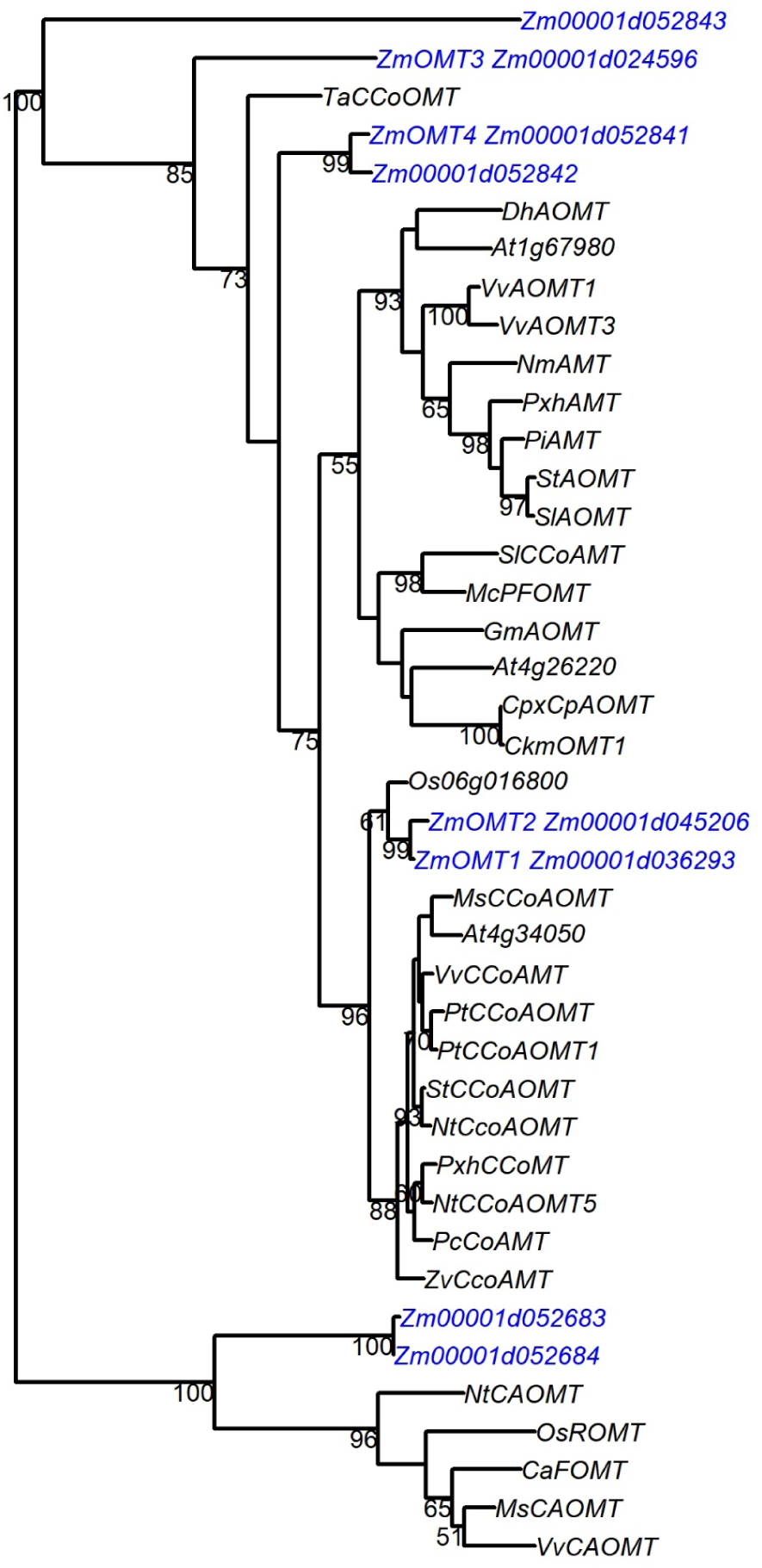
Phylogram of plant O-methyltransferases. Amino acid sequences were aligned and maximum likelihood tree was created. Branch numbers represent bootstrap values based on 100 replicates. Sequences correspond to table 4, and Zea mays sequences are highlighted in blue.

**Supplementary Figure 7:**
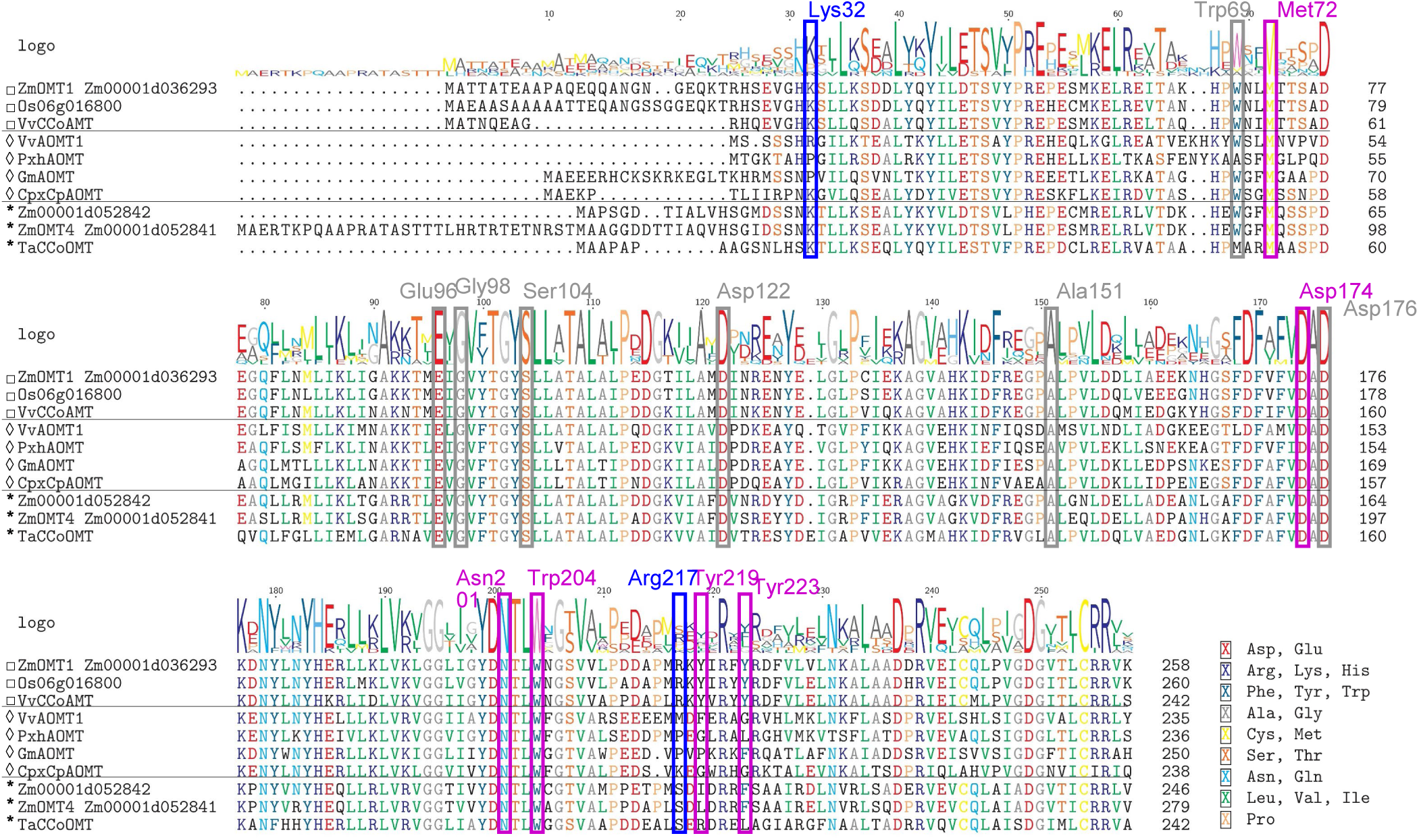
Alignment of Caffeoyl-CoA methyltransferases (□), Anthocyanin O-methyltransferases(◊), and candidate Zea Mays methyltransferases(*). Purple boxes and labels indicate Caffeoyl moiety binding, blue indicates CoA binding, and gray indicates SAM binding.

**Supplementary Figure 8:**
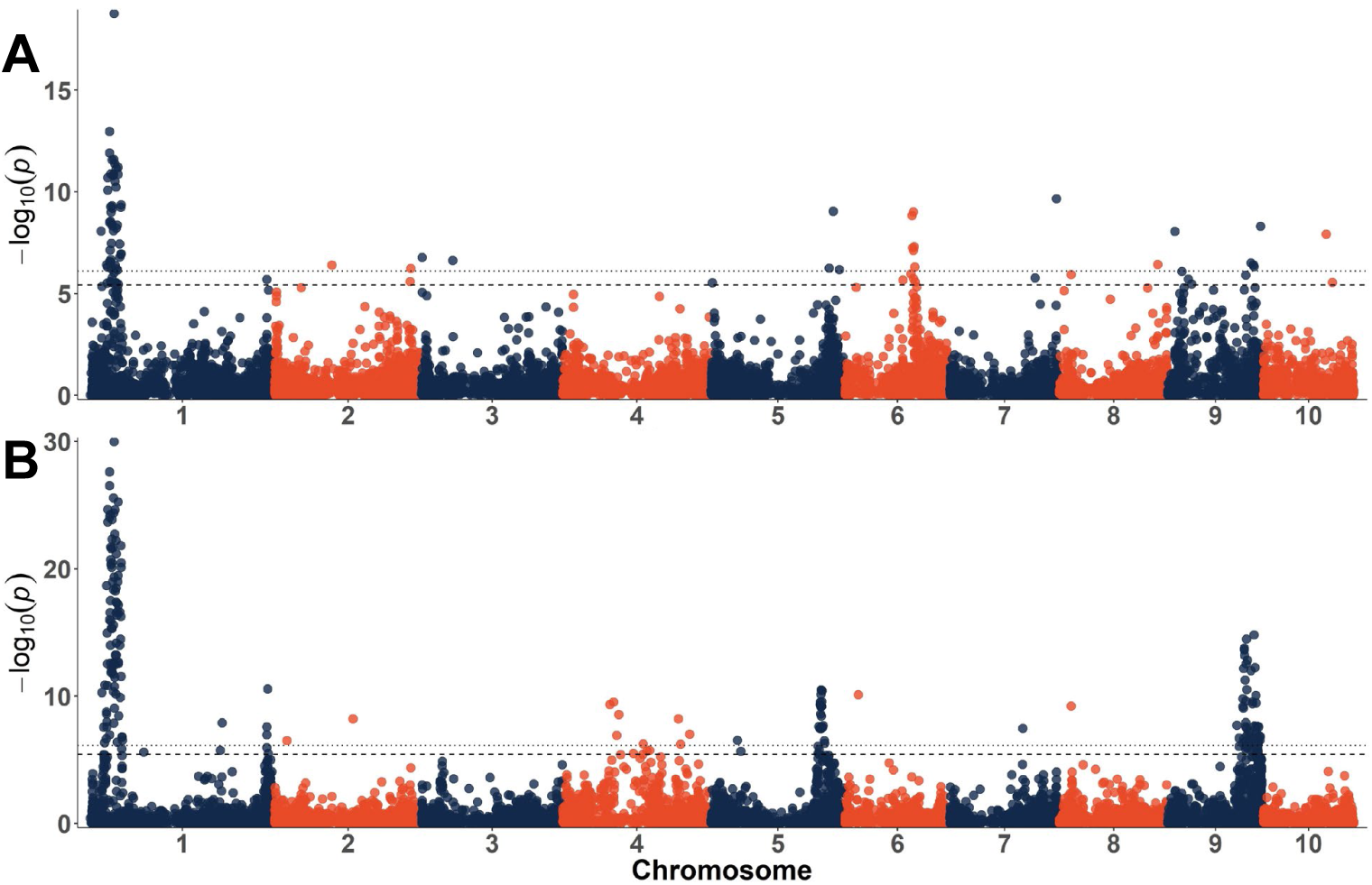
Manhattan plots for condensed forms using a flavone covariate (A) and flavones using a condensed forms covariate (B). Dotted lines indicate the experiment-wide significance thresholds using the Bonferroni correction. (α_E_=0.05 and α_E_=0.01)

**Supplementary Figure 9:**
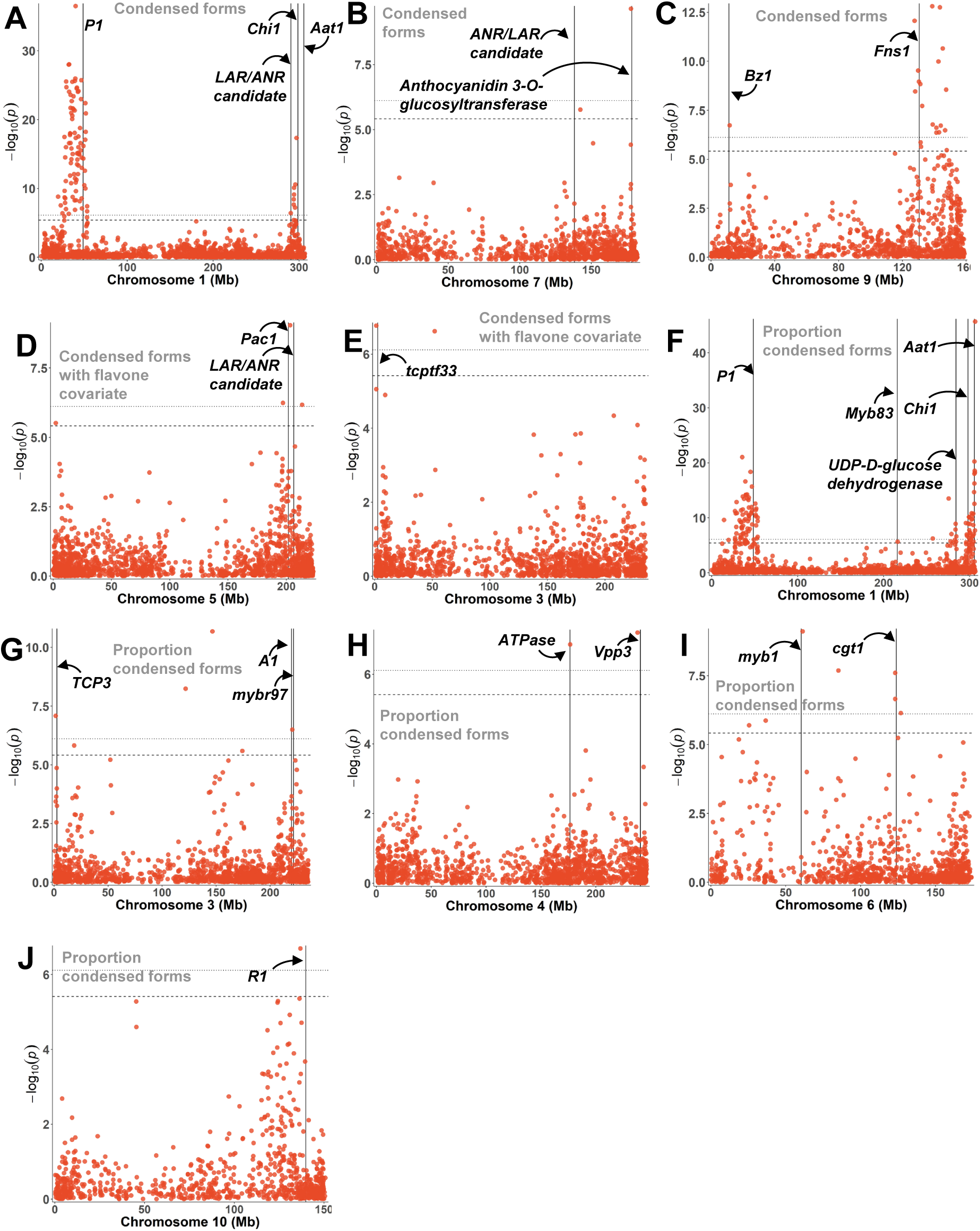
Individual chromosome plots for major signals identified for condensed forms (A-C), condensed forms with a flavone covariate (D-E), and the proportion of condensed forms compared to all anthocyanins (F-J). Chromosome numbers are listed in the x-axis label for each plot.

**Supplementary Figure 10:**
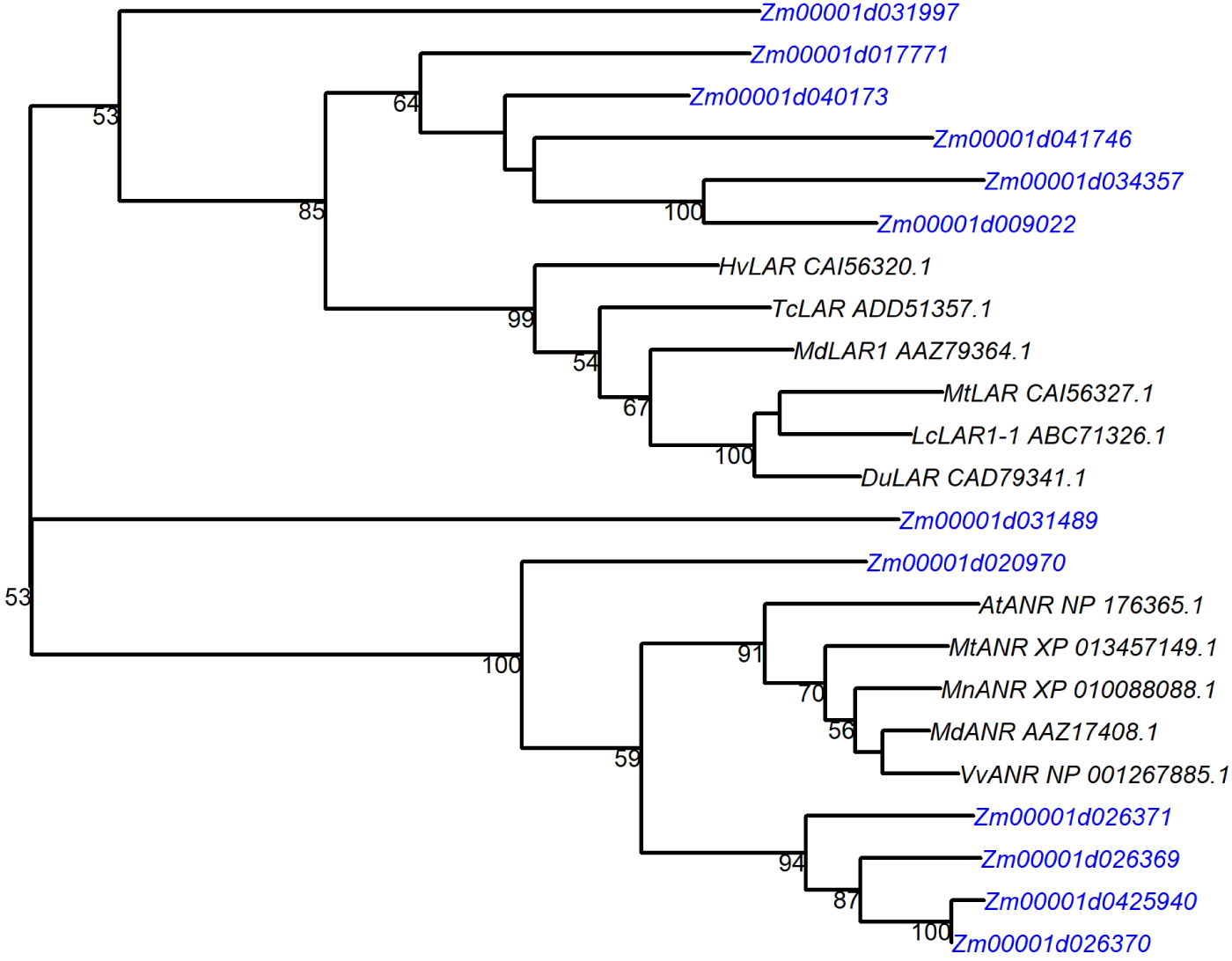
Maximum likelihood phylogeny based on peptide sequences of LAR and ANR sequences with possible LAR and ANR candidates. Branch numbers represent bootstrap values based on 100 replicates. GenBank proteins are listed with each LAR/ANR – *At, Arabidopsis thaliana; Mt, Medicago truncatula; Md, Malus domestica; Vv, Vitis vinifera; Mn, Morus notabilis; Lc, Lotus corniculatus; Tc, Theobroma cacao; Hv, Hordeum vulgare.*

**Supplementary Figure 11:**
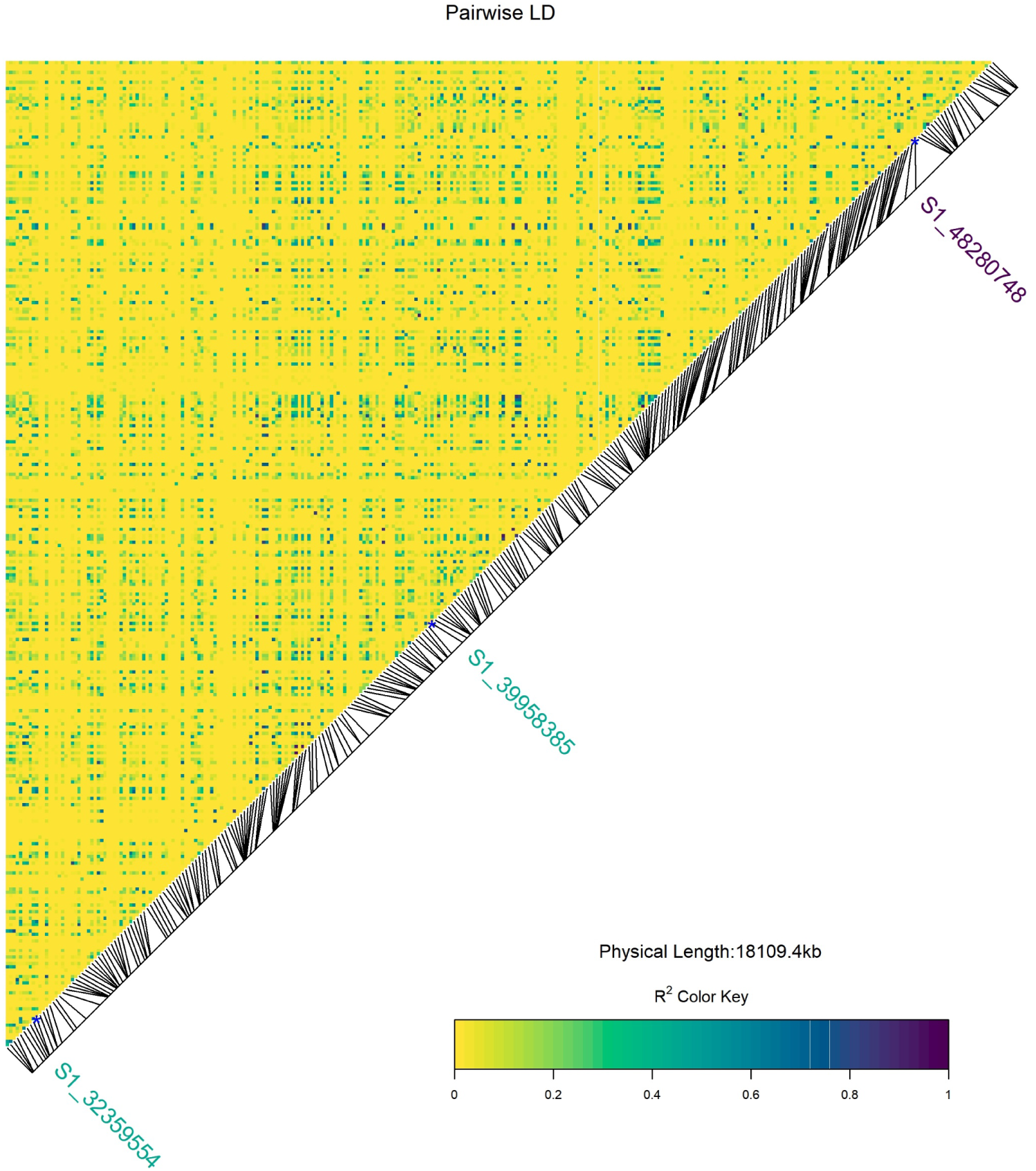
Pairwise LD for chromosome 1 from 33 to 50 Mb. SNPs labeled in green were highly significant SNPs from AR GWAS and the SNP labeled in dark purple is the SNP closest to *P1.*

**Supplementary Figure 12:**
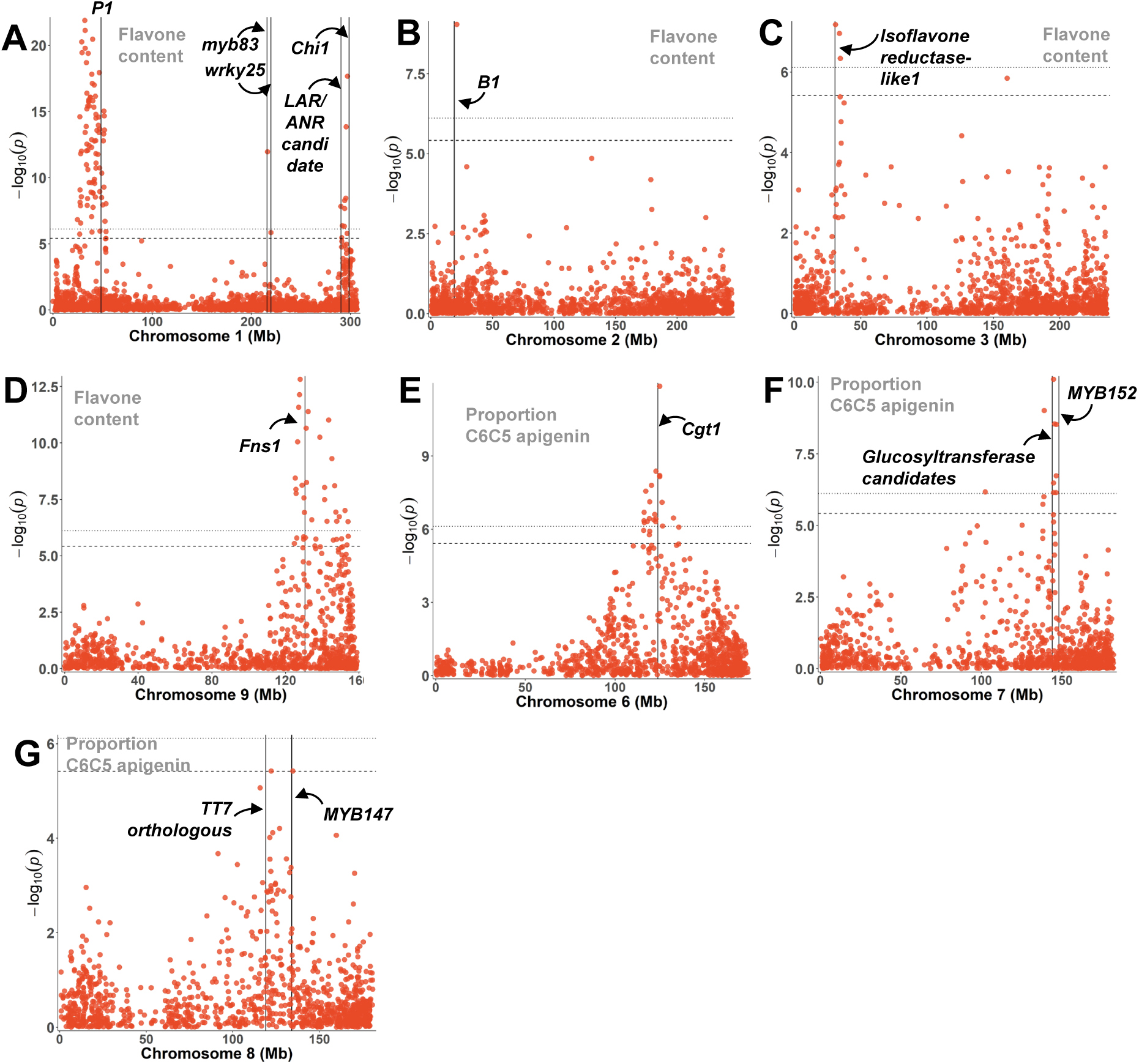
Individual chromosome plots for major signals identified for flavone content (A-D), and the proportion of *C* -hexosyl-*C* -pentosyl apigenin (E-G). Chromosome numbers are listed in the x-axis label for each plot.

